# Adaptive laboratory evolution rewires *Pseudomonas putida* for resource-efficient acetate assimilation

**DOI:** 10.64898/2026.08.20.746122

**Authors:** Nicolás Gurdo, Aparajitha Srinivasan, Tommaso Tagliani, Melanie Filbig, Nicolas T. Wirth, Josefin Johnsen, Garret W. O’Connell, Stefano Donati, Enrico Orsi, María V. G. Alván-Vargas, Yan Chen, Christopher J. Petzold, Matthew Blow, Thomas Eng, Till Tiso, Lars M. Blank, Adam Feist, Aindrila Mukhopadhyay, Pablo I. Nikel

## Abstract

Acetate is an attractive renewable two-carbon substrate for microbial biotechnology, but its toxicity limits growth and carbon-use efficiency at process-relevant concentrations. Here, we used adaptive laboratory evolution to improve acetate tolerance in a genome-reduced strain of *Pseudomonas putida* and combined whole-genome sequencing, reverse engineering, transcriptomics, proteomics, and ^13^C-acetate fluxomics to resolve the underlying adaptation mechanisms. Evolution under increasing acetate concentrations selected recurrent mutations in *gacA* and *fabB*, which encode a global response regulator and a fatty acid biosynthesis enzyme, respectively. Reverse engineering of these mutations recovered most of the evolved phenotype, including shorter lag phase and substantially higher biomass yield from acetate. Multi-omic analyses showed repression of type VI secretion systems, carbohydrate storage functions, fatty acid metabolism, and oxidative stress-associated proteins, indicating resource reallocation away from costly stress and non-essential programs. Fluxomics further revealed reduced EDEMP cycling and increased glyoxylate shunt flux, consistent with improved acetate-carbon retention in biomass. These results establish acetate tolerance in *P*. *putida* as a resource-efficiency phenotype and identify *gacA* and *fabB* as actionable targets for acetate-based bioproduction.

## 1. Introduction

Microbial bioproduction increasingly relies on alternative carbon sources to reduce feedstock costs and improve sustainability (1–4). Acetate is an attractive two-carbon (C_2_) substrate because it is inexpensive and can be generated through several renewable routes, including biomass hydrolysis, syngas fermentation, and microbial electrosynthesis (5–7). Acetate also constitutes a useful entry point into metabolism because it is directly converted into acetyl-coenzyme A (CoA) (8,9), a central precursor for growth and biosynthesis. However, acetate toxicity limits its use at process-relevant concentrations (10). As a weak organic acid, acetate can impair cytosolic pH homeostasis, disrupt central metabolism, increase maintenance demands, and reduce microbial fitness (11–15). A mechanistic understanding of acetate tolerance is therefore needed to exploit acetate more effectively in microbial cell factories.

*Pseudomonas putida* is a robust platform for bioproduction because it tolerates diverse environmental and chemical stresses (16–18), including organic acids (19). Genome-reduced variants of *P*. *putida* KT2440 have been developed as streamlined hosts for heterologous expression and metabolic engineering (20–22). Genome reduction can remove dispensable genetic elements and decrease the burden associated with non-essential functions (23–25), thereby improving genetic stability and freeing cellular resources for product formation (26). Overall, these properties make genome-reduced *P*. *putida* strains attractive backgrounds for examining acetate tolerance and for developing acetate-based bioproduction processes.

Acetate uptake in *P*. *putida* KT2440 occurs through the acetate permease ActP, followed by conversion into acetyl-CoA by acetyl-CoA synthetase (Acs). Acetyl-CoA can then enter the tricarboxylic acid (TCA) cycle, the glyoxylate shunt, lipid metabolism, or biosynthetic routes, depending on the physiological state of the cell. Previous engineering strategies have improved acetate utilization in Gram-negative bacteria by modifying endogenous metabolic pathways, including the glyoxylate shunt, or by targeting regulatory nodes across central metabolism, e.g., through modulation of isocitrate lyase activity (27–29). Recent work has also examined acetate use as a mixed carbon source with glucose in *P*. *putida* (30), and this C_2_ substrate is typically used as an energy source in selection schemes for synthetic one-carbon assimilation (31,32). Despite substantial progress, acetate toxicity remains a barrier to efficient growth and production, with rates, yields, and titers often remaining below values required for industrial implementation.

Adaptive laboratory evolution (ALE) offers a complementary route to improve substrate tolerance and uncover the genetic basis of complex phenotypes. ALE enriches beneficial mutations under defined selective pressures and has been widely used to improve growth, stress resistance, substrate uptake, and production traits (33–36). When combined with multi-omic analyses, ALE can also reveal how adaptation propagates across regulatory, metabolic, and physiological layers. Integrating whole-genome sequencing, transcriptomics, proteomics, and fluxomics enables a systems-level view of genotype-phenotype relationships and exposes the cellular strategies that support tolerance to specific stresses (37). A successful example of such framework is the adaptation of *Escherichia coli* to acetate as a carbon source. Sandberg et al. (38) evolved *E*. *coli* MG1655 on minimal medium containing acetate, nearly doubling the specific growth rate from to 0.43 h^−1^ and increasing acetate uptake from 12 to 20-23 mmol g cell dry weight (CDW) ^−1^ h^−1^.

Here, we applied ALE to a genome-reduced *P*. *putida* strain under gradually increasing acetate concentrations. We isolated acetate-tolerant clones, performed whole-genome sequencing to identify mutations associated with adaptation, reverse-engineered key mutations into the parental background, and integrated transcriptomics, proteomics, and ^13^C-based fluxomics to elucidate the mechanisms underlying acetate tolerance. This approach identified mutations in *gacA* and *fabB* as key contributors to the evolved phenotype and revealed systems-level changes in secretion systems, fatty acid metabolism, methionine and polyamine catabolism, iron uptake, and central carbon fluxes. Together, these analyses define genetic and metabolic targets for improving acetate assimilation in *P*. *putida* cell factories.

## 2. Materials and Methods

### 2.1. Bacterial strains, plasmids, and culture conditions

Strains and plasmids used in this study are listed in **Tables S1** and **S2**, respectively. *E*. *coli* DH5α λ*pir* was used as the cloning host for genetic manipulations. *E*. *coli* strains were cultivated at 37°C, and *P*. *putida* strains were cultivated at 30°C. Routine cloning, genome engineering, and strain propagation were performed in lysogeny broth (LB, 10 g L^−1^ tryptone, 5 g L^−1^ yeast extract, and 10 g L^−1^ NaCl). Liquid precultures were grown in 50-mL Falcon centrifuge tubes containing 10 mL of medium. Shaken-flask cultivations were performed in 250-mL baffled Erlenmeyer flasks containing 50 mL of medium. Liquid precultures were incubated at 250 rpm in a MaxQ 8000 incubator (Thermo Fisher Scientific Inc., Waltham, MA, USA), while shaken-flask cultures were incubated at 200 rpm in a New Brunswick Innova 42R shaker (Eppendorf SE, Hamburg, Germany). Solid media contained 15 g L^−1^ agar. Whenever required, kanamycin (Km) or gentamicin (Gm) was added at 50 μg mL^−1^ or 10 μg mL^−1^, respectively.

For ALE, clone screening, phenotypic characterization, and multi-omic experiments, strains were cultivated in minimal salt medium (MSM) (39) buffered with 5 g L^−1^ 3-(*N*-morpholino)propanesulfonic acid (MOPS). Potassium acetate was added at the concentrations indicated below, and the medium pH was adjusted to 7.0 after acetate addition. To inoculate experimental cultures, aliquots from precultures were harvested by centrifugation at 8,000×*g* for 5 min at room temperature, washed with MSM without carbon source, and resuspended in the final experimental medium to the desired starting optical density at 600 nm (OD_600_).

### 2.2 Adaptive laboratory evolution (ALE) on acetate

ALE of *P*. *putida* SEM1.4 was performed in 250-mL baffled Erlenmeyer flasks containing 50 mL of buffered MSM and supplemented with potassium acetate. Three parallel evolution experiments were initiated at 20 mM acetate, and acetate concentration was increased stepwise up to 180 mM. At each concentration, cultures were incubated at 30°C and 200 rpm until they reached mid-exponential phase (OD_600_ ∼ 0.5). Then, 500 μL of culture was transferred into fresh medium. In some cases, the transfer was done at the same acetate concentration for two additional passages before increasing acetate concentration further. The evolution experiment lasted 130 days, corresponding to ca. 200 passages, and was terminated when no further growth improvement was observed.

### 2.3 Screening of evolved populations and isolation of clones

The final evolved population was plated on MSM agar containing 180 mM acetate. Ten colonies with different colony sizes were selected, isolated, and stored at −80°C in cryopreservation solution containing 50% (v/v) glycerol, 0.1 M MgSO_4_, and 50 mM Tris·HCl (pH = 8). Phenotypic screening of selected clones was done in a Growth Profiler 960 system (Enzyscreen BV, Heemstede, The Netherlands) using square 96-well microtiter plates sealed with gas-permeable sandwich covers (Enzyscreen BV). MSM containing 5 g L^−1^ MOPS and acetate at 20, 60, 100, 140, or 180 mM was inoculated at an initial OD_600_ of 0.1 in a final volume of 300 μL per well. Bacterial growth was monitored every 15 min by optical image scanning in the Growth Profiler 960 apparatus. G-values, corresponding to integrated green values obtained from each scan and well, were transformed into equivalent OD_600_ values using calibration curves fitted to a Monod function (**Equation 1**):

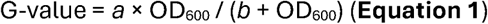

The parameters *a* and *b* were determined by non-linear regression. For calibration, the OD_600_ of serial culture dilutions in MSM was measured in a UV-1600PC spectrophotometer (VWR Inc., Radnor, PA, USA), and the corresponding images were acquired in the Growth Profiler 960 apparatus.

### 2.4 Whole-genome sequencing

Genomic DNA was extracted from cultures using the PureLink Genomic DNA Mini Kit (Thermo Fisher Scientific Inc., Waltham, MA, USA). Sequencing libraries were prepared using PlexWell (seqWell Inc., Beverly, MA, USA), and sequencing was performed on a NextSeq 500 instrument (Illumina Inc., San Diego, CA, USA) with an Illumina NextSeq mid-output kit using 300 cycles. Sequencing files were analyzed with an in-house pipeline described previously (40), based on Bowtie2 (41), using the *P*. *putida* KT2440 reference genome (42) with GenBank accession number AE015451. Average sequencing coverage was ca. 60× for clonal and population samples. For population samples, mutations with frequencies < 0.5 were excluded from the analysis (43) to filter out likely artifacts and focus on mutations enriched during ALE.

### 2.5 Reverse-engineering of mutations in P. putida SEM1.4

Point mutations identified during ALE were introduced into the parental strain by homologous recombination-mediated gene replacement. Uracil-excision (USER) cloning (44) was used to construct all plasmids. USER oligonucleotide primers containing the targeted single-nucleotide substitutions were designed with AMUSER (45) and are listed in **Table S3**. Phusion *U* Hot Start DNA Polymerase (Thermo Fisher Scientific Inc.) was used for PCR amplification, mutagenesis, and construction of USER-compatible DNA fragments. PCR products were analyzed on 1% (w/v) agarose gels and purified with the NucleoSpin Gel and PCR Clean-up kit (Macherey-Nagel GmbH, Düren, Germany). Purified amplicons were assembled into plasmid pSNW2 (21) using USER enzyme (New England BioLabs Inc., Ipswich, MA, USA) according to the manufacturer’s instructions, and the resulting plasmids were transformed into *E*. coli DH5α λ*pir* by heat shock (46). Gene replacement was performed using I-*SceI*-mediated recombination (47) with inducible self-curing vectors (21). Positive clones, initially screened by green fluorescence and colony PCR, were grown in LB. Cultures were washed with 0.3 M sucrose and transformed by electroporation with the self-curing helper plasmid pQURE6·H (22). Cells were recovered overnight in LB containing 5 mM 3-methylbenzoate and then streaked onto LB agar containing Gm and 5 mM 3-methylbenzoate. Gene substitution events were screened by red fluorescence, colony PCR, and DNA sequencing. Single colonies were then grown in LB without selection to cure the helper plasmid. Non-fluorescent colonies were selected, and plasmid loss was verified by colony PCR, parallel plating on LB or LB and Gm, and DNA sequencing. Confirmed mutants were grown in LB and preserved as cryostocks.

### 2.6 Phenotypic characterization of reverse-engineered strains

The parental strain *P*. *putida* SEM1.4, the acetate-evolved strain SEM1.4Evo, and the reverse-engineered double mutant SEM1.4 GacA^E197K^ FabB^L77P^, together with other control strains indicated in the text, were phenotypically characterized in 250-mL baffled Erlenmeyer flasks containing 50 mL of MSM supplemented with 100 mM acetate and buffered with 5 g L^−1^ MOPS. This acetate concentration was selected because it was the lowest concentration that clearly differentiated the strains in microplate experiments. Cultures were inoculated at an initial OD_600_ of 0.1 and incubated at 30°C and 200 rpm. OD_600_ was measured during cultivation, and acetate consumption was quantified by high-performance liquid chromatography (HPLC). Samples were centrifuged at 10,000×*g* for 5 min at 4°C, and supernatants were stored at −20°C until HPLC analysis. Cell dry weight (CDW) was determined under the same cultivation conditions. At different cultivation times, six 10-mL samples were collected per strain, corresponding to two samples per biological replicate, in pre-weighed 15-mL Falcon tubes. OD_600_ was measured before sample processing. Biomass was harvested by centrifugation at 8,000×*g* for 10 min at 4°C and washed twice with MSM without carbon source. Tubes containing the biomass were frozen at −20°C and lyophilized under vacuum until constant weight. Growth parameters were derived from these measurements using the QurvE software (48).

### 2.7 Metabolite analysis

Acetate consumption was analyzed using a Dionex UltiMate 3000 HPLC system (Thermo Fisher Scientific Inc.) equipped with an Aminex HPX-87H ion-exclusion column (300×7.8 mm, Bio-Rad Laboratories Ltd., Hercules, CA, USA), a refractive index detector, and UV detectors set to 210, 260, 277, and 304 nm (49). The column was maintained at 45°C, and samples were eluted with 5 mM H_2_SO_4_ at a constant flow rate of 0.6 mL min^−1^. HPLC data were processed with the Chromeleon 7.1.3 software (Thermo Fisher Scientific Inc.). Acetate concentrations were calculated from peak areas using calibration curves prepared with six standard concentrations ranging from 0 to 250 mM.

### 2.8 Transcriptomic analysis

Cells growing on MSM supplemented with 100 mM acetate were harvested in exponential phase at OD_600_ = 0.6. Culture samples of 1.5 mL were centrifuged at 12,000×*g* for 2 min. Supernatants were removed, and RNA was isolated from cell pellets using the GeneJET RNA Purification Kit according to the manufacturer’s instructions (Thermo Fisher Scientific Inc.). RNA quality and concentration were determined using a Qubit RNA High Sensitivity Assay Kit on a Qubit fluorometer (Thermo Fisher Scientific Inc.). Three biological replicates were processed per condition.

For RNA-Seq library generation and sequencing, ribosomal RNA was depleted from 100 ng total RNA using QIAseq FastSelect 5S/16S/23S, rRNA Plant, rRNA Yeast, and custom rRNA Algae depletion kits (Qiagen NV, Hilden, Germany). Libraries were prepared using the TruSeq Stranded mRNA kit (Illumina Inc.). Heat-fragmented RNA fragments of 300-400 bp were reverse-transcribed into first-strand cDNA using random hexamers and SuperScript II Reverse Transcriptase (Thermo Fisher Scientific Inc.), followed by second-strand synthesis. Double-stranded cDNA fragments were processed by A-tailing, ligated to NEXTFLEX UDI barcodes (PerkinElmer Inc., Waltham, MA, USA), and enriched using 10 cycles of PCR. Prepared libraries were quantified using the KAPA Biosystems next-generation sequencing library qPCR kit (Roche AG, Basel, Switzerland) on a LightCycler 480 real-time PCR instrument (Roche AG). Sequencing was performed on an Illumina NovaSeq instrument using NovaSeq XP v1.5 reagent kits and an S4 flow cell with a 2×151 indexed run configuration.

The raw FASTQ reads were filtered and trimmed using the JGI QC pipeline, yielding filtered FASTQ files. BBDuk was used to identify artifact sequences by k-mer matching with k = 25, allowing one mismatch, and detected artifacts were trimmed from the 3′ end of reads. RNA spike-in reads, PhiX reads, and reads containing ambiguous bases were removed. Quality trimming was performed using the Phred trimming method set at Q6. After trimming, reads below the length threshold were removed. The minimum length threshold was set to 25 bp or one third of the original read length, whichever was longer. Filtered reads from each library were aligned to the reference genome using HISAT2 v2.2.1 with the -k 1 flag (50). Strand-specific coverage bigWig files were generated using deepTools v3.1 (51). Raw gene counts were generated from gff3 annotations using featureCounts (52). Only primary hits assigned to the reverse strand were included in raw gene counts using the -s 2, -p, and -primary options. Raw gene counts were used to evaluate correlations among biological replicates using Pearson correlation and to determine which replicates were retained for differential gene expression analysis.

DESeq2 v1.30.0 (53) was used to identify differentially expressed genes between pairs of conditions. Genes with adjusted *p*-values < 0.05 were considered differentially expressed (54). A complete list of differentially expressed genes is provided in the Supplementary Data (**Data S1**) and at the JGI Genome Portal under Project ID 1445921.

### 2.9 Proteomic analysis

*P*. *putida* SEM1.4 and its derivative strains detailed in the text were precultured overnight in MSM supplemented with 20 mM acetate. Experimental cultures were performed in shaken flasks containing MSM supplemented with 100 mM acetate and inoculated at an initial OD_600_ of 0.1. Four biological replicates were collected in mid-exponential phase at OD_600_ = 0.6. Cells were harvested by centrifugation at 10,000×*g* for 5 min at 4°C. Supernatants were removed, and cell pellets were frozen at −80°C until further processing.

Protein extraction from cell pellets and tryptic peptide preparation followed an established proteomic sample preparation protocol. Cell pellets were resuspended in Qiagen P2 lysis buffer (Qiagen NV). Proteins were precipitated by addition of 1 mM NaCl and four volumes of acetone, followed by two washes with 80% (v/v) acetone in water. The recovered protein pellet was homogenized in 100 mM ammonium bicarbonate in 20% (v/v) methanol. Protein concentration was determined using the DC Protein Assay (Bio-Rad Laboratories Ltd.). Proteins were reduced with 5 mM *tris*(2-carboxyethyl)phosphine for 30 min and alkylated with 10 mM iodoacetamide for 30 min at room temperature in the dark. Proteins were digested overnight with trypsin at a 1:50 trypsin-to-total-protein ratio.

Peptide samples were analyzed on an Agilent 1290 ultra-high-performance liquid chromatography system (Agilent Technologies Inc., Santa Clara, CA, USA) coupled to an Orbitrap Exploris 480 mass spectrometer (Thermo Fisher Scientific Inc.). Peptides were loaded onto an Ascentis ES-C18 column (Sigma-Aldrich Co., St. Louis, MO, USA) and eluted using a 10-min gradient from 98% solvent A and 2% solvent B to 65% solvent A and 35% solvent B. Solvent A was 0.1% (v/v) formic acid in water, and solvent B was 0.1% (v/v) formic acid in acetonitrile. Eluting peptides were introduced into the mass spectrometer operating in positive-ion mode and analyzed by data-independent acquisition. The acquisition method used a duty cycle of three survey scans from *m*/*z* = 380 to *m*/*z* = 985 and 45 MS2 scans with a precursor isolation width of 13.5 *m*/*z* to cover the mass range. DIA raw data files were analyzed with DIA-NN.

The database used for DIA-NN analysis in library-free mode consisted of the latest *P*. *putida* KT2440 UniProt proteome FASTA sequences (55) plus common proteomic contaminants. DIA-NN automatically determined mass tolerances based on first-pass analysis of the samples and optimized mass accuracy. Retention time extraction windows were determined individually for each MS run using the automated optimization procedure implemented in DIA-NN. Protein inference was enabled, and the quantification strategy was set to Robust LC = High Accuracy. DIA-NN reports were filtered using a global false discovery rate of 0.01 at both precursor and protein-group levels.

Protein abundance was estimated using the Top3 approach, which calculates the average MS signal intensity of the three most abundant tryptic peptides detected for each protein (56–58). Proteomics data were analyzed using a customized R script in RStudio v1.3.1093. Only proteins detected in all samples were retained. Abundance values were normalized by quantile normalization using the R package qsmooth. Statistical differences between groups were assessed using Student’s *t*-test. Data were log_2_-transformed for fold-change comparisons, and volcano plots were generated using VolcaNoseR (59). Proteins with an absolute log_2_(fold change) > 1.5 and −log_10_(adjusted *p*-value) > 1.3 were considered differentially abundant.

### 2.10 Fluxomic analysis

Precultures were inoculated from freshly plated colonies of *P*. *putida* SEM1.4, SEM1.4Evo, and SEM1.4 GacA^E197K^ FabB^L77P^ in MSM supplemented with 20 mM unlabeled acetate and 5 g L^−1^ MOPS. Before inoculation of experimental cultures, cells were harvested by centrifugation at 10,000×*g* for 5 min and washed twice with MSM without carbon source. Cells were then inoculated individually at an initial OD_600_ = 0.02 in 100-mL shaken flasks containing 20 mL of MSM supplemented with 100 mM labeled acetate. The labeling experiments used 99% [1-^13^C]-sodium acetate, 99% [2-^13^C]-sodium acetate, or 50% [U-^13^C_2_]-sodium acetate. The inoculum carried into labeled cultures was kept below 1% (v/v) of the final sampled biomass concentration to minimize interference from unlabeled inoculum in subsequent flux calculations (60). All experiments were performed with three biological replicates and two technical replicates. Samples were harvested when cultures reached OD_600_ = 2 for analysis of proteinogenic amino acids and cellular sugars. Samples were centrifuged at 10,000×*g* for 5 min. Supernatants were removed, and pellets were frozen at −80°C until further processing. Cell pellets were thawed on ice and resuspended in 200 μL of 6 M HCl. Samples were incubated at 105°C for 16-24 h to hydrolyze biomass (61). Hydrolyzed samples were filtered using a 96-well filter plate (MultiScreenHTS HV Filter Plate, 0.45 μm, hydrophilic, clear, non-sterile, EMD Millipore Corp., Burlington, MA, USA) by centrifugation at 1,500×*g* for 2 min. Filtered samples were freeze-dried and stored at −80°C. Dried samples were derivatized in two steps. First, hydrolysates were resuspended in 50 μL dimethylformamide. Second, this solution was transferred into glass vials containing 50 μL *N*-*tert*-butyldimethylsilyl-*N*-methyltrifluoroacetamide with 1% (w/w) *tert*-butyldimethylchlorosilane and incubated at 85°C for 1 h. Derivatized samples were aliquoted into glass vials with inserts for GC-MS analysis within 12 h of derivatization.

Samples were injected into an Agilent 7890A GC-MS system (Agilent Technologies Inc.) equipped with an Agilent HP-5ms capillary column with a length of 30 m, inner diameter of 0.25 mm, and film thickness of 0.25 μm. Samples were measured in full-scan mode using the following temperature program: 120°C for 1 min, ramp to 160°C at 4°C min^−1^ and hold for 5 min, ramp to 270°C at 4°C min^−1^ and hold for 3 min, ramp to 310°C at 20°C min^−1^ and hold for 1 min, and ramp to 120°C at 60°C min^−1^.

For analysis of cellular sugar monomers, glucose and glucosamine, cell pellets were hydrolyzed in 250 μL of 2 M HCl for 2 h at 100°C (62). Cell debris was removed by filtration using a 96-well filter plate. Hydrolysates were freeze-dried and stored at −80°C. Dried residues were derivatized with 2% (w/v) methoxylamine in pyridine at 80°C (1 h) and silylated with *N*,*O*-*bis*(trimethylsilyl)trifluoroacetamide (Macherey-Nagel GmbH & Co., Düren, Germany) at 80°C (30 min). Derivatized analytes were quantified on an Agilent 7890A GC-MS system (Agilent Technologies Inc.) equipped with an HP-5ms capillary column. Samples were measured in full-scan mode using a temperature program from 120°C to 300°C. Raw chromatographic data from amino acid and sugar analyses were integrated using SmartPeak (63) and corrected for natural isotope abundance in derivatization agents using INCA (64). Mass isotopomer distributions from 14 proteinogenic amino acids were used for flux estimation, excluding cysteine and tryptophan because they degrade during hydrolysis (65,66). Glutamate and aspartate were also used as proxies for glutamine and asparagine, respectively, due to deamination during protein hydrolysis. For sugar analysis, glucose fragments at *m*/*z* = 319 and 554 and glucosamine fragments at *m*/*z* = 319 and 553 were included as described by Kohlstedt and Wittmann (67).

The acetate-dependent metabolic network for *P*. *putida* SEM1.4 was built from the genome-scale model *i*JN1462 (68) with the modifications indicated by Bujdoš et al. (69). The reconstruction included 72 reactions from central carbon metabolism; a list of reactions and the cognate carbon atom transitions are provided in **Table S4**. Fluxes were estimated with INCA using specific growth rates and acetate uptake or secretion rates as constraints, and the biomass equation was taken from Czajka et al. (70). Relative intracellular fluxes were inferred by minimizing the weighted sum of squared residuals between simulated and experimental labeling data. Flux estimation was repeated at least 20 times with random initial values until convergence. Goodness of fit was assessed using a χ^2^ test, and 95% confidence intervals were calculated from the sensitivity of the residuals to flux-parameter variation (71). Flux distributions were visualized by mapping computed values onto custom metabolic maps using the R package fluctuator.

### 2.11 STRING interaction network analysis and enrichment analysis

Interaction networks for differentially regulated transcripts and proteins were obtained using the STRING v12.0 database (72). The input lists contained genes or proteins identified as differentially regulated or differentially abundant, submitted by gene names or UniProt identifiers. A default confidence score threshold of 0.4, corresponding to medium confidence, was used to filter interactions. Interaction networks were visualized and clustered according to functional association (73). Networks were further analyzed to identify highly connected genes or proteins and biological pathways enriched in each dataset.

### 2.12 AlphaFold prediction and structural characterization

The three-dimensional structure of GacA was predicted with AlphaFold 2.0 using default settings (74). The five highest-ranking predictions were analyzed and validated using the local distance difference test, SPServer (75), and ProSA-Web (76). Visualization, structural superimposition, and in silico mutagenesis were performed with PyMOL v2.3.4 (Schrödinger Inc., New York, NY, USA). Original protein sequences were obtained from the *Pseudomonas* Database (77). Homolog searches and sequence analyses were performed using the AlphaFold template-search algorithm, NCBI BLASTP, InterPro (78), and Jalview v2.11 (79).

### 2.13 Data and statistical analysis

All experiments reported were independently repeated at least three times, and the mean values of the corresponding parameters ± standard deviations are presented. In some cases, statistical significance between groups was assessed using an unpaired Student’s *t*-test. Levels of significance, when relevant, were identified with asterisk symbols (*) as follows: * *p* < 0.05, ** *p* < 0.01, and *** *p* < 0.001. The **Data S1** file lists RNA-Seq, proteomic, and fluxomic data comparisons across the strains used in this study. Data analysis was performed with MS Excel (Microsoft Corp., Redmond, WA, USA) and Prism 8 (GraphPad Software Inc., San Diego, CA, USA) unless otherwise specified.

## 3. Results and Discussion

### 3.1 ALE enhances acetate tolerance in *P. putida*

*P*. *putida* can use a broad range of carbon substrates (18), including acetate, a weak C_2_ organic acid that becomes inhibitory at high concentrations (80). Acetic acid, the protonated form, can diffuse across the cell envelope and dissociate in the cytosol, perturbing pH homeostasis, increasing energetic burden, and triggering pleiotropic physiological damage (12,81). Acetate tolerance is therefore expected to depend on coordinated responses across transport, central metabolism, redox balancing, membrane physiology, and stress-response systems, as reported for other Gram-negative species (82–84).

To investigate the mechanisms underlying acetate tolerance, we implemented an ALE strategy. Strain SEM1.4, a genome-reduced strain (55,85) derived from *P*. *putida* EM42 (86), was evolved in MSM under gradually increasing acetate concentrations, from 20 to 180 mM. Cultures were transferred once they reached mid-exponential phase, which under these conditions corresponds to an OD_600_ ∼ 0.5. During the initial stages, from 20 to 80 mM acetate, transfers occurred every 6-12 h. At higher concentrations, transfers were performed daily or after more than 24 h, as the time required to reach the target OD_600_ was longer. The evolution experiment lasted 130 days, corresponding to ca. 200 passages, and was terminated when no further growth improvement was observed after a 12-day incubation in the presence of 180 mM acetate. Under this final condition, cultures reached only OD_600_ values of ca. 0.5-0.6.

The evolved population from the final passage, hereafter referred to as *P*. *putida* SEM1.4Evo, was plated on MSM agar containing 180 mM acetate. Ten colonies were isolated from this population to capture phenotypic variation while maintaining strong selective pressure (**Figure 1A**). These clones were then phenotypically characterized under increasing acetate concentrations. Whole-genome sequencing was used to identify mutations that arose during evolution, and selected mutations were introduced into the parental background by reverse engineering. The resulting strains were analyzed using transcriptomics, proteomics, and ^13^C-metabolic flux analysis to connect genotype, physiology, and systems-level responses to acetate stress (**Figure 1B**).

**Figure 1.**
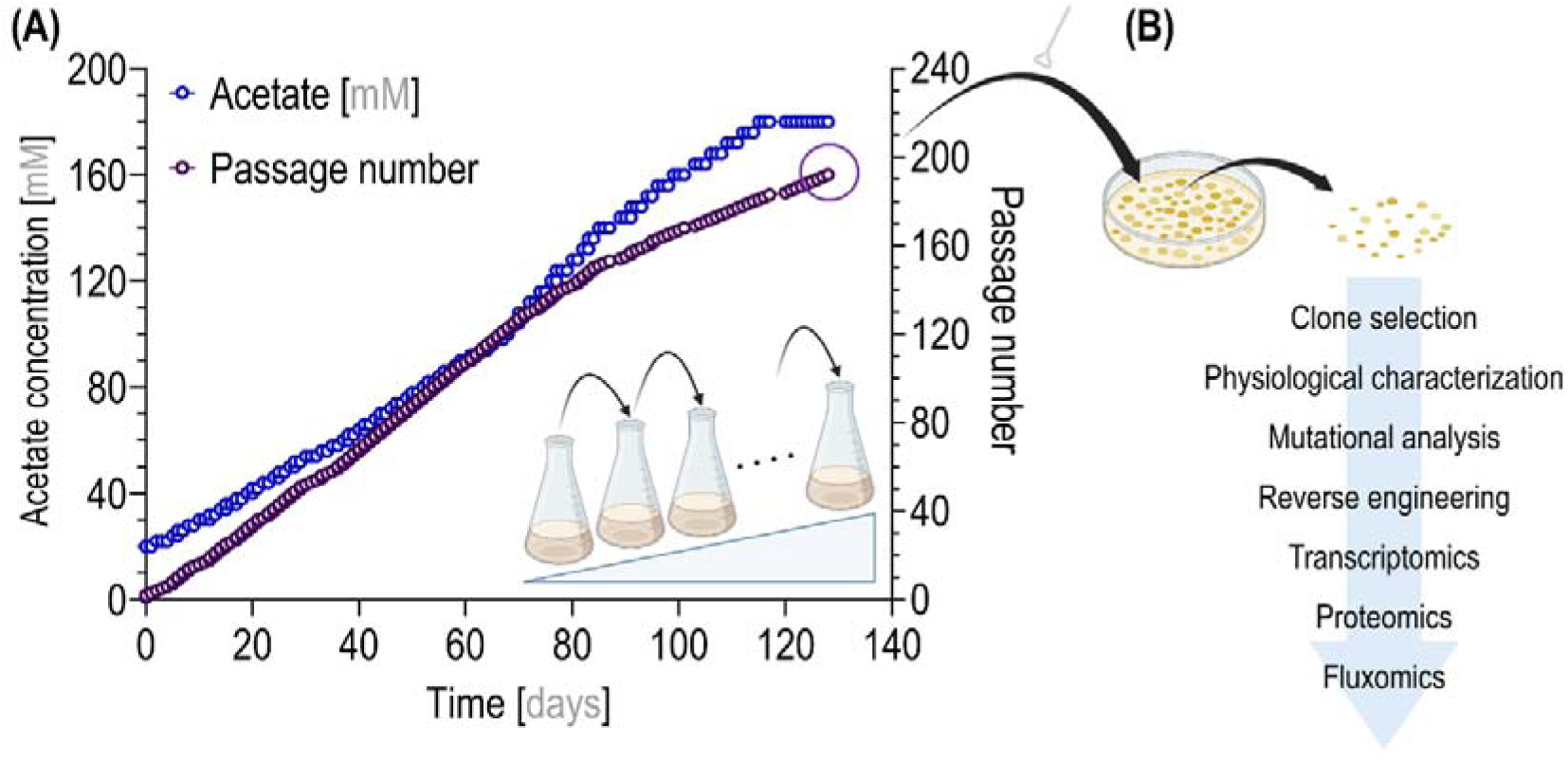
Workflow for investigating acetate toxicity in *P. putida* SEM1.4. **(A)** Adaptive laboratory evolution (ALE) strategy to increase tolerance to high concentrations of acetate. **(B)** Physiological and multi-omic characterization of selected evolved clones.

### 3.2 Growth profiling of acetate-evolved clones reveals improved performance under high acetate concentrations

The ten isolated clones and the parental strain *P*. *putida* SEM1.4 were challenged with increasing acetate concentrations in MSM. Growth was monitored for 48 h in medium containing 20, 60, 100, 140, or 180 mM acetate. At 20 mM acetate, the parental strain and evolved clones displayed similar growth profiles, although most evolved clones reached higher final OD_600_ values after ca. 10 h of cultivation (**Figure 2A**). Differences became more pronounced as acetate concentration increased. At 60 and 100 mM acetate, most evolved clones displayed shorter lag phases than the parental strain, with reductions of ca. 25% and 50%, respectively (**Figure 2B** and **2C**).

**Figure 2.**
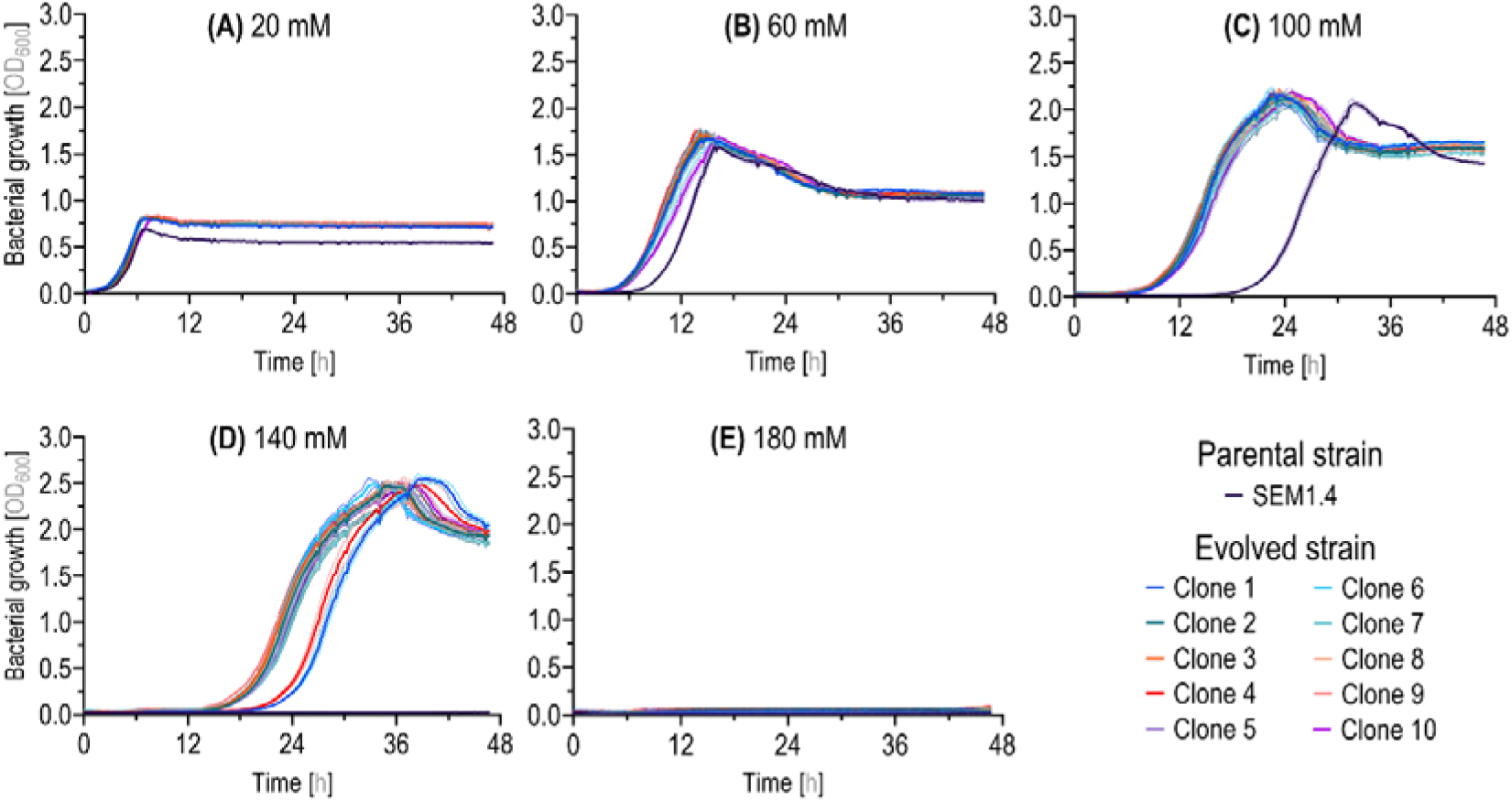
Acetate tolerance of selected evolved clones and the parental strain in 96-well microtiter plates. Growth profiles of ten clones and the parental strain in minimal salt medium (MSM) containing 5 g L^−1^ MOPS and supplemented with acetate at **(A)** 20 mM, **(B)** 60 mM, **(C)** 100 mM, **(D)** 140 mM, and **(E)** 180 mM. Solid lines represent mean optical density at 600 nm (OD_600_), and shaded areas indicate the standard deviation from four biological replicates.

At 140 mM acetate, the parental strain did not show measurable growth after 48 h, whereas the evolved clones reached OD_600_ values of ca. 2.5 (**Figure 2D**). This phenotype indicates that ALE substantially increased the ability of *P*. *putida* SEM1.4 to initiate and sustain growth under acetate concentrations that strongly inhibit the parental background. At 180 mM acetate, no detectable biomass formation was observed for either the parental strain or the evolved clones, indicating that this concentration exceeded the tolerance range achieved under the conditions used here (**Figure 2E**). Overall, growth profiling showed that acetate-driven evolution mainly reduced lag phase and extended the concentration range in which *P*. *putida* SEM1.4 could grow, rather than enabling growth at the highest acetate concentration imposed during the evolution experiment. This phenomenon has also been observed in *P*. *putida* KT2440 evolved for tolerance to *p*-coumaric acid (87).

### 3.3 Mutational analysis identifies recurrent mutations in global regulatory and fatty acid metabolism genes

To identify mutations associated with acetate tolerance, the ten selected clones were analyzed by whole-genome sequencing. The analysis focused on mutations located within coding sequences and present in most clones. Four recurrent nonsynonymous mutations were identified in genes associated with regulatory control, fatty acid biosynthesis, and nucleoid organization: *gacA* (*PP_4099*, formerly known as *uvrY*), encoding the response regulator of the GacS/GacA two-component system; *fabB* (*PP_4175*), encoding 3-oxoacyl-ACP (acyl-carrier-protein) synthase I; *PP_0371*, encoding a LysR-family transcriptional regulator; and *hupB* (*PP_2303*), encoding the β subunit of a DNA-binding protein (**Table 1**).

**Table 1.**
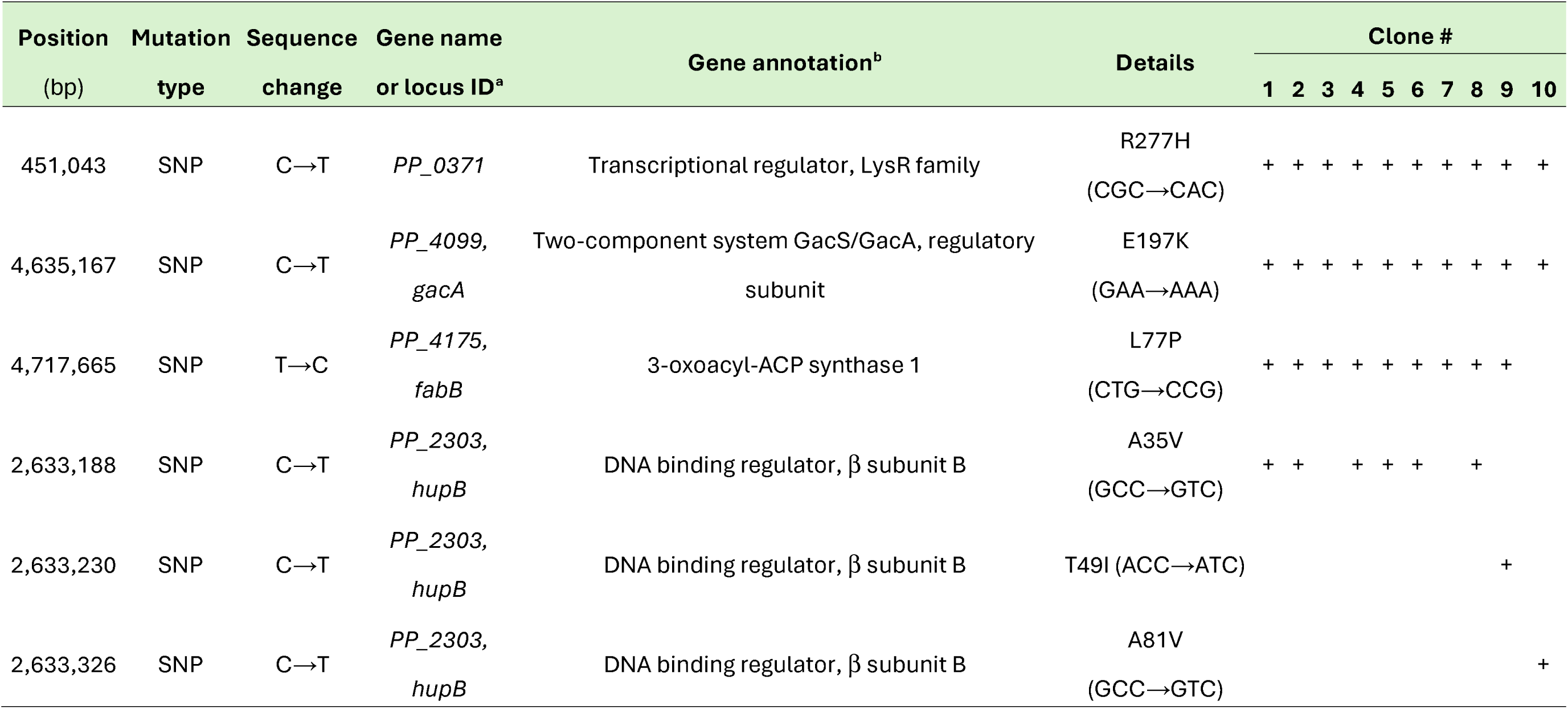
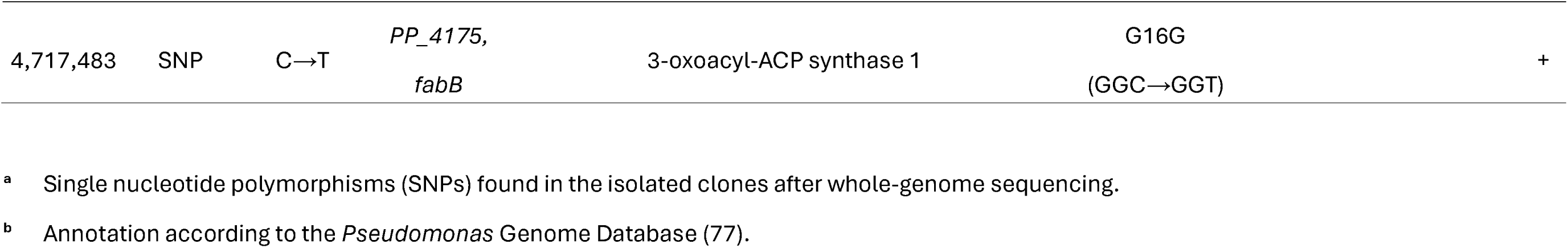
Mutational analysis of selected clones after the evolution on acetate.^a^.

All ten clones carried mutations in *gacA* and *PP_0371*, nine clones carried a mutation in *fabB*, and six clones carried the *hupB***^A35V^** mutation (**Table 1**). Additional *hupB* and *fabB* variants appeared in single clones, suggesting lower prevalence within the evolved population. The high recurrence of the *gacA***^E197K^**, *PP_0371***^R277H^**, and *fabB***^L77P^** mutations indicates strong selection for changes affecting global regulation and envelope-associated metabolism during growth under acetate stress.

The GacS/GacA two-component system controls broad adaptive programs in pseudomonads, including central carbon metabolism, secondary metabolism, quorum sensing, motility, and biofilm formation (88,89). Disruption of *gacA* has also been linked to improved fitness and bioproduction phenotypes in other *P*. *putida* backgrounds (90). The recurrence of *gacA* mutations in all acetate-evolved clones therefore pointed to attenuation of this regulatory system as a potential route to acetate tolerance. In parallel, the repeated *fabB* mutation suggested that membrane lipid metabolism also contributed to adaptation. Consistent with this observation, mutations affecting both *gacS* and *gacA* were independently identified during acetate-driven ALE of *P*. *putida* KT2440, and deletion of either gene improved growth on acetate (91). FabB catalyzes a key elongation step in fatty acid biosynthesis and contributes to membrane composition, fluidity, and integrity in *Pseudomonas* species (92–94). Mutations in *hupB* and *PP_0371* further suggested that acetate tolerance involved broader regulatory and nucleoid-associated changes. Overall, the mutational spectrum indicates that acetate tolerance in *P*. *putida* SEM1.4 is a multifactorial trait shaped by coordinated changes in global regulation, fatty acid metabolism, and genome-associated transcriptional control.

Loss-of-function mutations in the GacS/GacA two-component system are a recurrent outcome of ALE in *P*. *putida* KT2440, and the *gacA* mutations recovered here during evolution on acetate follow this pattern. The *gacS* gene was mutated in all tolerance-ALE and ALE lineages evolved on the hydroxycinnamates *p*-coumaric and ferulic acid (87), while *fleQ*, *gacA*/*gacS*, *wbpL*, and *flgE* were among the most frequently mutated loci during evolution on the lignin-derived aromatic vanillate (95). More recently, convergent loss-of-function mutations in *gacS* under all conditions and in *gacA* under terephthalate and terephthalate/ethylene glycol conditions emerged during evolution on PET-derived substrates. Reverse engineering showed that deletion of *gacA* or *gacS* shortened lag phase and improved growth and substrate utilization (96). Across these studies, disruption of GacS/GacA has been interpreted as a largely substrate-independent adaptation to well-mixed, nutrient-defined batch cultivation, where downregulation of costly flagellar, biofilm, and secondary-metabolic programs redirects resources toward growth. This interpretation echoes the long-recognized tendency of *Pseudomonas* to accumulate spontaneous *gacS*/*gacA* mutants in laboratory cultures, first characterized in biocontrol strains where these mutations abolish secondary-metabolite production and where loss of costly secondary metabolism can drive mutant accumulation (97–100). The emergence of *gacA* mutations under acetate selection therefore extends this convergent GacS/GacA signature to a non-aromatic carbon source.

### 3.4 Reverse engineering of recurrent mutations links gacA and fabB variants to acetate tolerance

To evaluate the contribution of recurrent ALE mutations to acetate tolerance, we introduced selected single-nucleotide substitutions into the genome of the parental strain *P*. *putida* SEM1.4. Mutation selection was based on prevalence across the evolved clones, using a threshold > 50% (**Table 1**). This criterion prioritized four nonsynonymous substitutions (nucleotide changes in the corresponding gene are indicated in bold, underlined): PP_0371^R277H^ (CGC→C**<u>A</u>**C), GacA^E197K^ (GAA→**<u>A</u>**AA), FabB^L77P^ (CTG→C**<u>C</u>**G), and HupB^A35V^ (GCC→G**<u>T</u>**C). The corresponding single mutants were constructed by scarless allelic exchange, and their growth was evaluated under increasing acetate concentrations (**Figure 3**).

**Figure 3.**
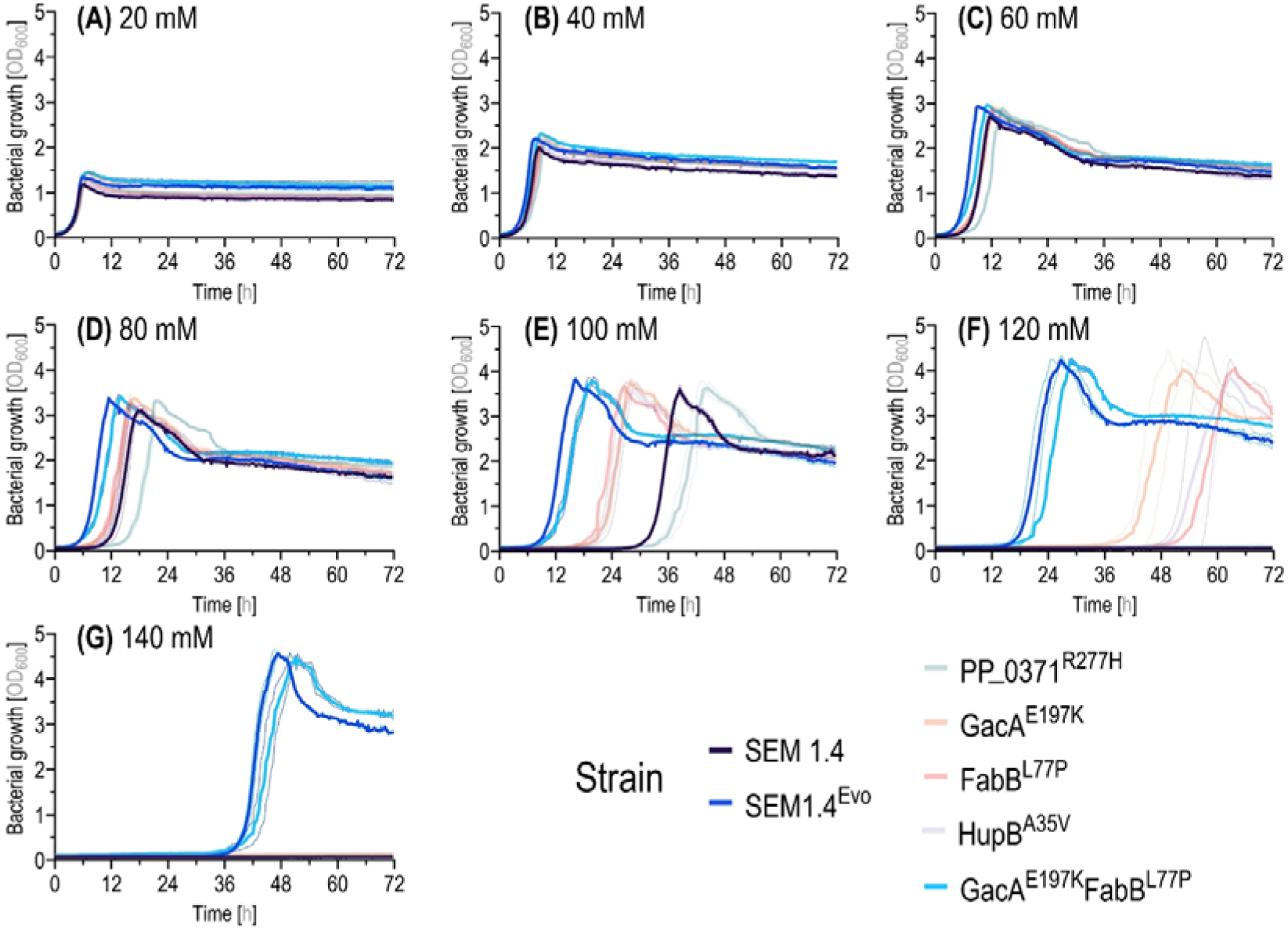
Evaluation of parental, evolved, and reverse-engineered strains carrying key single-nucleotide polymorphisms (SNPs) at high acetate concentrations. Reverse-engineered mutants PP_0371^R277H^, GacA^E197K^, FabB^L77P^, HupB^A35V^, and GacA^E197K^ FabB^L77P^ were tested in minimal salt medium (MSM) supplemented with acetate at **(A)** 20 mM, **(B)** 40 mM, **(C)** 60 mM, **(D)** 80 mM, **(E)** 100 mM, **(F)** 120 mM, and (G) 140 mM, and compared to the parental P. putida SEM1.4 strain and its evolved derivative (SEM1.4Evo). Solid lines represent mean values, and shaded areas indicate the standard deviation from four biological replicates.

At 20 mM acetate, all strains displayed comparable growth profiles (**Figure 3A**). Differences became evident at higher concentrations. Strains carrying *gacA***^E197K^**, *fabB***^L77P^**, or *hupB***^A35V^** showed shorter lag phases than the parental strain, particularly above 60 mM acetate (**Figure 3B-E** and **Table 2**). The strongest single-mutation effects were observed for the reverse-engineered *gacA***^E197K^** and *fabB***^L77P^** variants. In contrast, strain SEM1.4 PP_0371^R277H^ showed impaired growth across the tested conditions and was not pursued further. The parental strain failed to grow at 120-140 mM acetate after 72 h, whereas several reverse-engineered strains retained measurable growth at 120 mM acetate and the GacA^E197K^ FabB^L77P^ double mutant grew at 140 mM acetate (**Figure 3F** and **3G**).

**Table 2.**
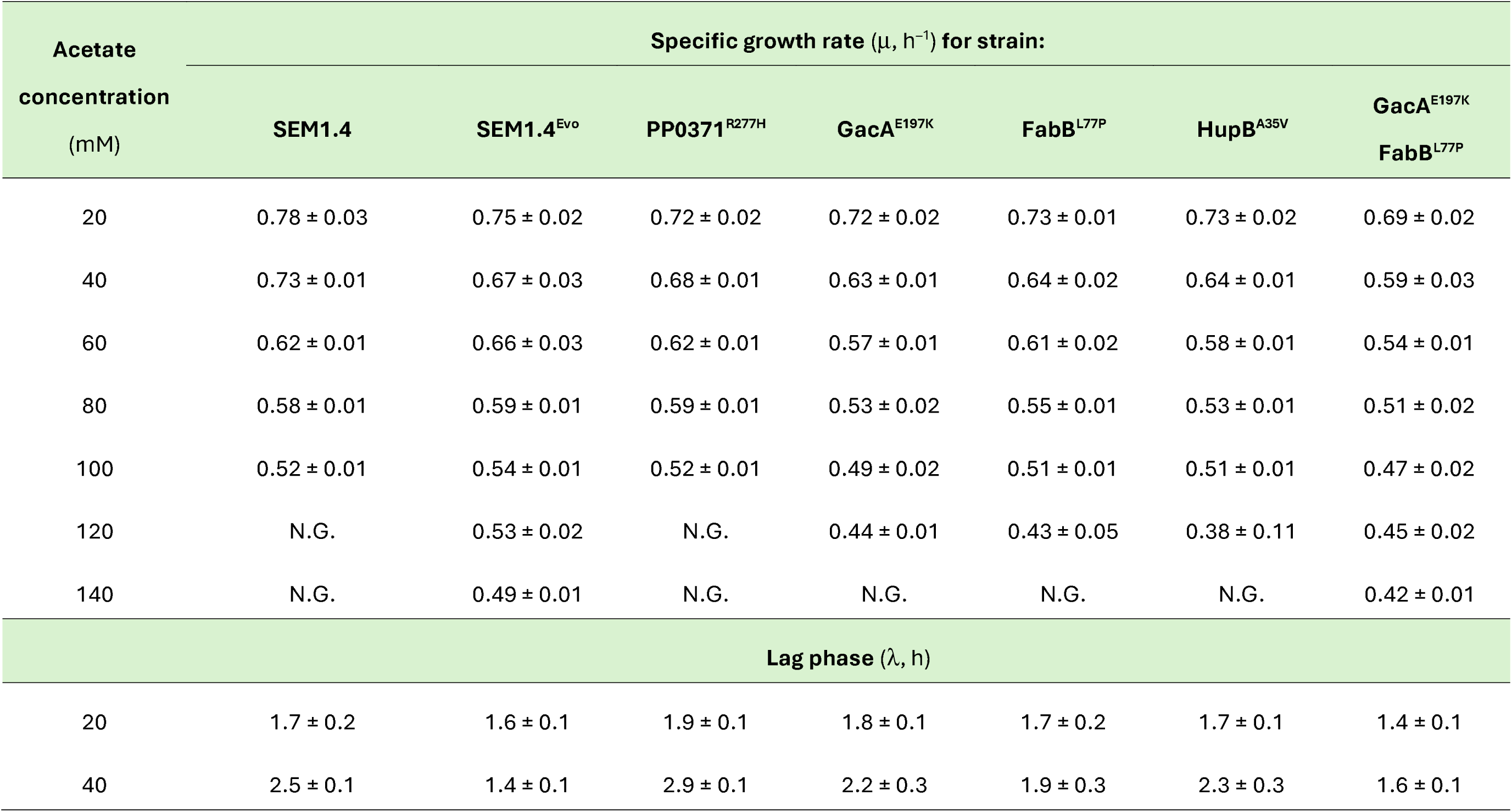
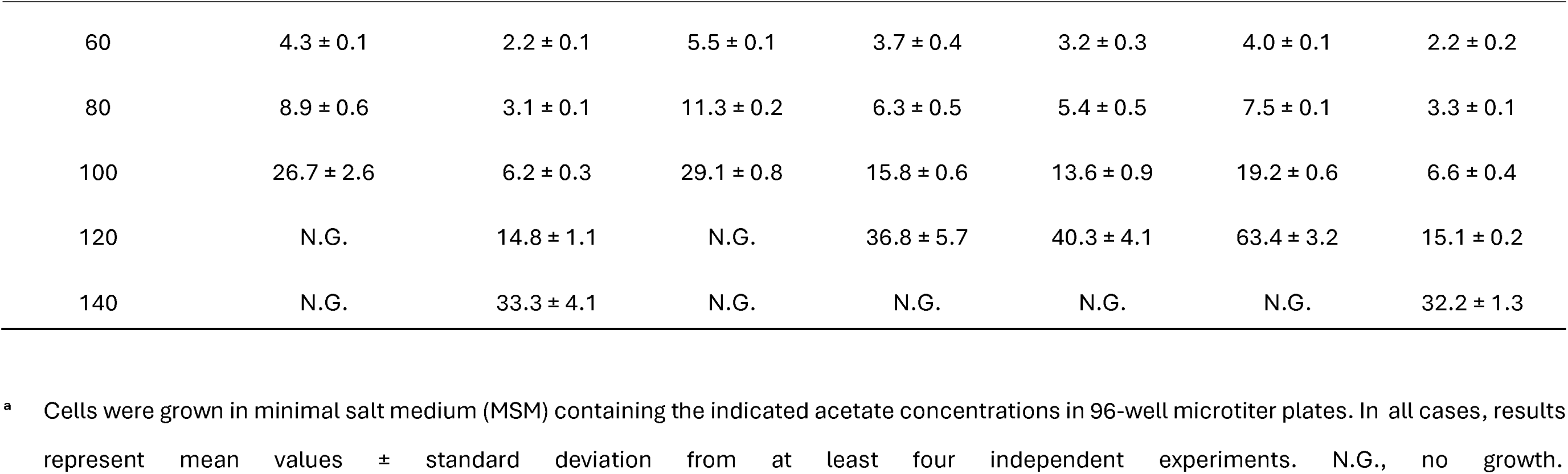
Growth parameters^a^ of P. putida SEM1.4, SEM1.4^Evo^, and reverse-engineered strains grown with increasing acetate concentrations.

| Acetate<br>concentration<br>(mM) | Specific growth rate ($\mu$, h<sup>-1</sup>) for strain: | | | | | | |
| --- | --- | --- | --- | --- | --- | --- | --- |
|  | SEM1.4 | SEM1.4<sup>Evo</sup> | PP0371<sup>R277H</sup> | GacA<sup>E197K</sup> | FabB<sup>L77P</sup> | HupB<sup>A35V</sup> | GacA<sup>E197K</sup><br>FabB<sup>L77P</sup> |
| 20 | 0.78 ± 0.03 | 0.75 ± 0.02 | 0.72 ± 0.02 | 0.72 ± 0.02 | 0.73 ± 0.01 | 0.73 ± 0.02 | 0.69 ± 0.02 |
| 40 | 0.73 ± 0.01 | 0.67 ± 0.03 | 0.68 ± 0.01 | 0.63 ± 0.01 | 0.64 ± 0.02 | 0.64 ± 0.01 | 0.59 ± 0.03 |
| 60 | 0.62 ± 0.01 | 0.66 ± 0.03 | 0.62 ± 0.01 | 0.57 ± 0.01 | 0.61 ± 0.02 | 0.58 ± 0.01 | 0.54 ± 0.01 |
| 80 | 0.58 ± 0.01 | 0.59 ± 0.01 | 0.59 ± 0.01 | 0.53 ± 0.02 | 0.55 ± 0.01 | 0.53 ± 0.01 | 0.51 ± 0.02 |
| 100 | 0.52 ± 0.01 | 0.54 ± 0.01 | 0.52 ± 0.01 | 0.49 ± 0.02 | 0.51 ± 0.01 | 0.51 ± 0.01 | 0.47 ± 0.02 |
| 120 | N.G. | 0.53 ± 0.02 | N.G. | 0.44 ± 0.01 | 0.43 ± 0.05 | 0.38 ± 0.11 | 0.45 ± 0.02 |
| 140 | N.G. | 0.49 ± 0.01 | N.G. | N.G. | N.G. | N.G. | 0.42 ± 0.01 |
| Lag phase ($\lambda$, h) | | | | | | | |
| 20 | 1.7 ± 0.2 | 1.6 ± 0.1 | 1.9 ± 0.1 | 1.8 ± 0.1 | 1.7 ± 0.2 | 1.7 ± 0.1 | 1.4 ± 0.1 |
| 40 | 2.5 ± 0.1 | 1.4 ± 0.1 | 2.9 ± 0.1 | 2.2 ± 0.3 | 1.9 ± 0.3 | 2.3 ± 0.3 | 1.6 ± 0.1 |
| 60 | 4.3 ± 0.1 | 2.2 ± 0.1 | 5.5 ± 0.1 | 3.7 ± 0.4 | 3.2 ± 0.3 | 4.0 ± 0.1 | 2.2 ± 0.2 |
| 80 | 8.9 ± 0.6 | 3.1 ± 0.1 | 11.3 ± 0.2 | 6.3 ± 0.5 | 5.4 ± 0.5 | 7.5 ± 0.1 | 3.3 ± 0.1 |
| 100 | 26.7 ± 2.6 | 6.2 ± 0.3 | 29.1 ± 0.8 | 15.8 ± 0.6 | 13.6 ± 0.9 | 19.2 ± 0.6 | 6.6 ± 0.4 |
| 120 | N.G. | 14.8 ± 1.1 | N.G. | 36.8 ± 5.7 | 40.3 ± 4.1 | 63.4 ± 3.2 | 15.1 ± 0.2 |
| 140 | N.G. | 33.3 ± 4.1 | N.G. | N.G. | N.G. | N.G. | 32.2 ± 1.3 |

<sup>a</sup> Cells were grown in minimal salt medium (MSM) containing the indicated acetate concentrations in 96-well microtiter plates. In all cases, results represent mean values ± standard deviation from at least four independent experiments. N.G., no growth.

Because the strains carrying the *gacA***^E197K^** and *fabB***^L77P^** alleles had the clearest improvements among the single mutants, we combined both mutations in strain *P*. putida SEM1.4 GacA^E197K^ FabB^L77P^ to examine whether they jointly reproduced the evolved phenotype. The double mutant closely matched strain SEM1.4Evo across most acetate concentrations, with a markedly reduced lag phase and sustained growth at acetate concentrations that inhibited the parental strain (**Figure 3** and **Table 2**). These results indicate that the *gacA***^E197K^** and *fabB***^L77P^** mutations recapitulate much of the acetate-tolerant phenotype obtained through ALE, although additional mutations or population-level effects may contribute to the full behavior of the evolved strain.

### 3.5 Physiological characterization of the reverse-engineered double mutant substantiates the evolved phenotype and links acetate tolerance to resource-efficient carbon use

To evaluate whether the reverse-engineered mutations reproduced the evolved phenotype under standard shaken-flask conditions, we characterized *P*. *putida* SEM1.4, strain SEM1.4Evo, and the double mutant strain SEM1.4 GacA^E197K^ FabB^L77P^ in MSM containing 100 mM acetate. This concentration was selected because it was the lowest acetate level that clearly differentiated the strains during microplate screening (**Figure 3E**). Sampling times for subsequent multi-omic analyses corresponded to mid-exponential phase.

Shaken-flask cultivations confirmed that acetate-driven evolution improved growth under acetate stress. The parental strain displayed a lag phase of ca. 22 h, whereas strain SEM1.4Evo and the double mutant had shorter lag phases of ca. 9 and 15 h, respectively (**Figure 4A**). Acetate consumption followed the growth profiles, although the parental strain had the highest specific acetate uptake rate (*q*_A_), reaching 34.1 ± 4.7 mmol g_CDW_^−1^ h^−1^, substantially higher than 25.8 ± 0.8 mmol g_CDW_^−1^ h^−1^ for strain SEM1.4Evo and 23.8 ± 1.6 mmol g_CDW_^−1^ h^−1^ for the double mutant (**Figure 4B** and **4C**). In this context, faster acetate uptake by the parental strain did not lead to higher biomass formation. This phenotype indicates that acetate tolerance was not driven by faster acetate consumption alone, but by more efficient conversion of acetate carbon into biomass.

**Figure 4.**
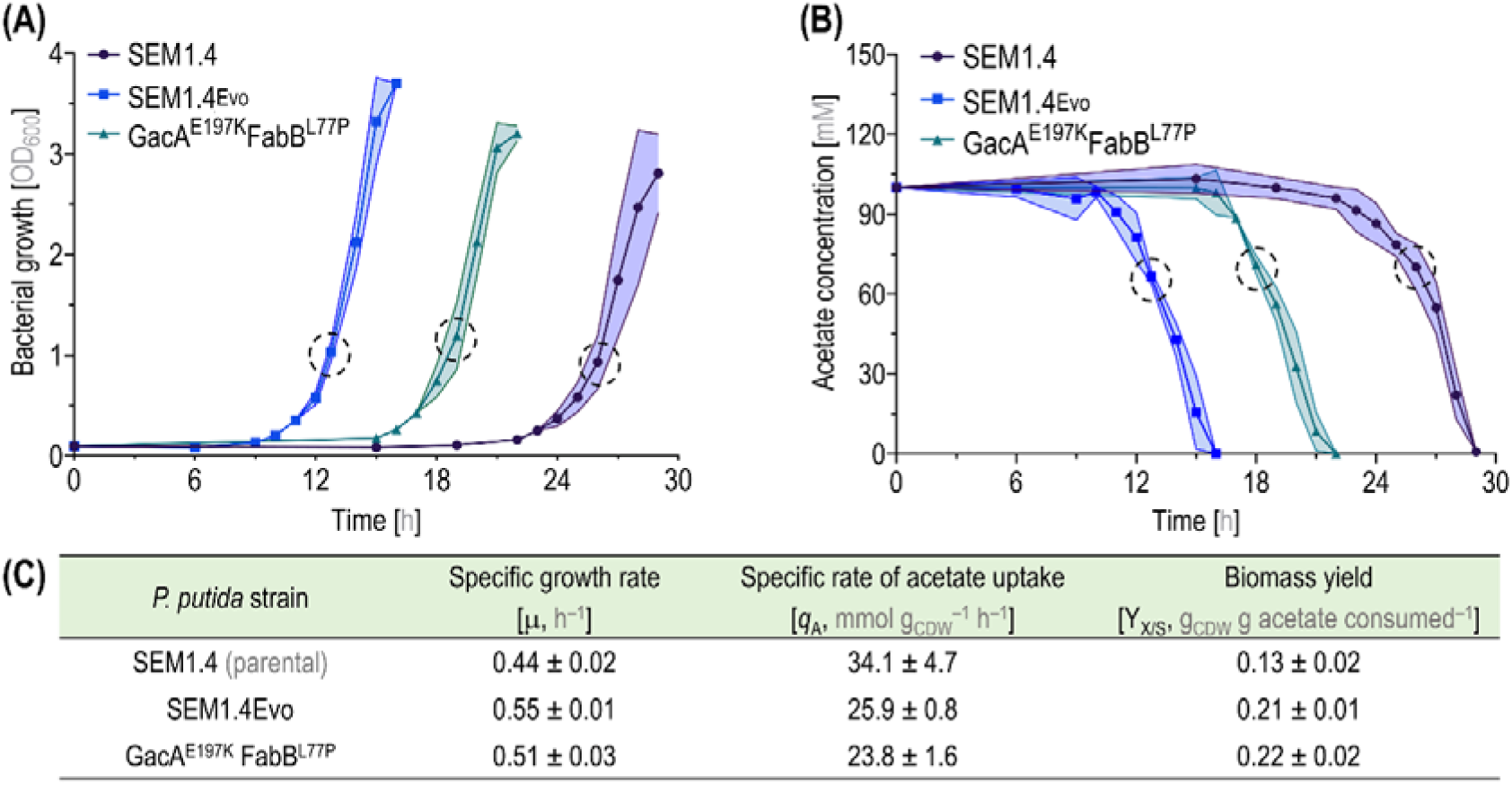
Physiological characterization in shaken-flask cultivations. *P. putida* SEM1.4, SEM1.4Evo, and the reverse-engineered GacA^E197K^ FabB^L77P^ strain were grown in MSM supplemented with 100 mM acetate to evaluate **(A)** growth curves and **(B)** acetate consumption. Solid lines represent mean values, and shaded areas indicate the standard deviation from three biological replicates; dotted circles indicate the mid-exponential growth phase. Specific growth rates, acetate uptake rates, and biomass yields are shown in **(C)**, where each parameter is shown as the mean ± standard deviation. CDW, cell dry weight.

This yield phenotype is central to the mechanistic interpretation of the evolved state. Both strain SEM1.4Evo and the double mutant showed ca. 20% higher specific growth rates and 60% higher biomass yields than the parental strain (**Figure 4C**). Higher biomass yield means that a larger fraction of consumed acetate was retained in cellular material rather than being lost through CO_2_ release, maintenance metabolism, oxidative-stress management, or futile cycling through central metabolism. The double mutant reproduced most of this improvement, linking the *gacA***^E197K^** and *fabB***^L77P^** mutations to a resource-efficient acetate-use phenotype.

These physiological differences defined the basis for the multi-omic analyses that follow. If the evolved and double mutant strains use acetate more efficiently, their transcriptomes, proteomes, and fluxomes should reveal reduced investment in costly non-essential functions and a reorganization of central carbon metabolism toward carbon conservation. The sections below test this model and show that the higher biomass yield is associated with global regulatory rewiring, reduced expression of energetically expensive stress and secretion programs, altered fatty acid metabolism, and flux redistribution toward acetate-conserving routes.

### 3.6 Transcriptomic profiling links acetate tolerance to repression of costly cellular programs

To identify transcriptional changes associated with acetate tolerance, RNA-Seq analysis was performed with the parental strain SEM1.4, strain SEM1.4Evo, and the reverse-engineered double mutant strain SEM1.4 GacA^E197K^ FabB^L77P^ during growth in MSM containing 100 mM acetate. Samples were collected in exponential phase, and differential gene expression was evaluated across three pairwise comparisons: strain SEM1.4Evo *versus* the parental strain SEM1.4, the double mutant *versus* the parental strain SEM1.4, and the double mutant *versus* strain SEM1.4Evo (**Figure 5A-D**). These comparisons were designed to determine whether the two reverse-engineered mutations reproduced the regulatory state selected during acetate-driven evolution.

**Figure 5.**
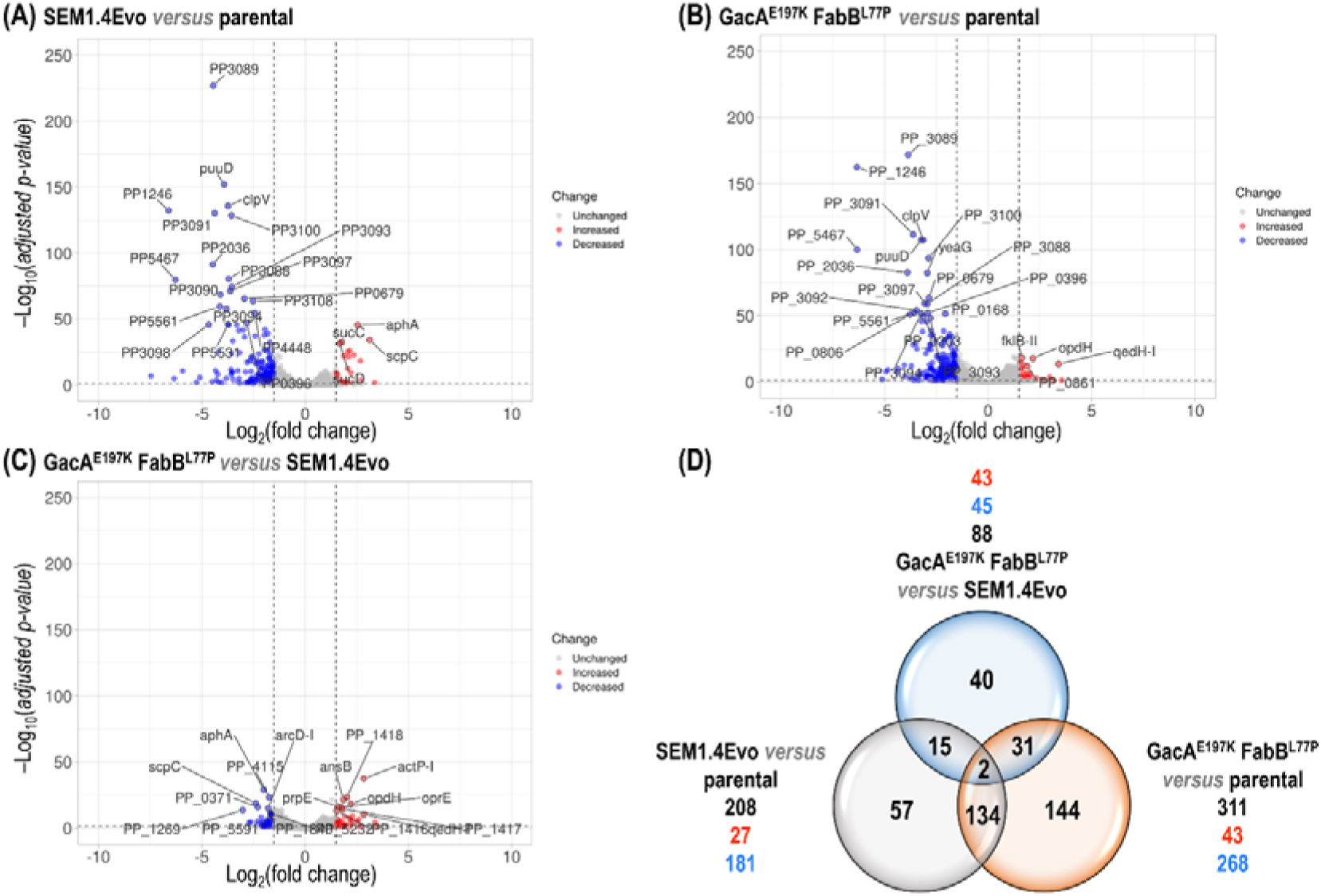
Transcriptomic analysis. Volcano plots for RNA-Seq data showing differentially expressed transcripts in *P. putida* SEM1.4Evo versus the parental strain (SEM1.4) **(A)**, reverse-engineered GacA^E197K^ FabB^L77P^ versus the parental strain **(B)**, and reverse-engineered GacA^E197K^ FabB^L77P^ versus SEM1.4Evo **(C)**. Three biological replicates were grown in MSM supplemented with 100 mM acetate and harvested during mid-exponential phase for transcriptomic analysis. Horizontal thresholds were set at 1.3 with p < 0.05, and vertical thresholds at −1.5 and +1.5 log_2_(fold change). **(D)** Venn diagram showing significantly regulated transcripts shared among the comparisons in **(A–C)**. Red and blue numbers indicate upregulated and downregulated genes, respectively, whereas dark gray indicates the total number of significantly regulated transcripts in each comparison.

Relative to the parental strain SEM1.4, strain SEM1.4Evo showed broad transcriptional remodeling, with 208 differentially expressed genes, including 27 upregulated and 181 downregulated genes (**Figure 5A** and **5D**). The induced gene set included those encoding putrescine catabolism, amino acid transport, energy metabolism, and transcriptional regulation. These included *aphA*, encoding an acetylpolyamine aminohydrolase, *potF-IV*, encoding a putrescine-binding periplasmic protein, *gabD-II*, encoding an NADP^+^-dependent succinate-semialdehyde dehydrogenase, and the amino-acid transporter genes *PP_1298* and *PP_1299*. The upregulation of *scpC*, *sucC*, and *sucD* (**Figure S1A**) further pointed to increased use of succinyl-CoA–linked reactions and alternative routes feeding succinate metabolism. This response is consistent with a shift toward acetate-carbon assimilation routes that support growth while maintaining precursor supply under stress.

The most prominent transcriptional signature in strain SEM1.4Evo was gene repression. The repressed gene fraction was dominated by secretion systems, environmental sensing, carbohydrate storage, glycosyl transferase functions, and fatty acid biosynthesis. Several components of the type VI secretion system (T6SS) were repressed, including genes from the K1-T6SS and K3-T6SS clusters, orphan *hcp*-associated loci, and *vgrG*-linked clusters (**Figure S1B**). Since T6SS expression is energetically costly and controlled by the Gac/Rsm regulatory cascade in pseudomonads, repression of this program suggests a mechanistic link between the *gacA***^E197K^** mutation and resource reallocation under acetate stress. The evolved strain also downregulated two-component regulatory genes, glycogen-, trehalose-, and maltose-associated genes, and a fatty acid biosynthesis cluster spanning *PP_3775* to *PP_3787*. These changes suggest reduced investment in non-essential secretion, sensing, storage, and envelope-building functions during growth on acetate.

The reverse-engineered double mutant showed a transcriptional response that largely mirrored the evolved strain. Relative to the parental strain SEM1.4, strain SEM1.4 GacA^E197K^ FabB^L77P^ displayed extensive differential gene expression dominated by downregulated genes (**Figure 5B** and **5D**). Upregulated functions again included polyamine transport and catabolism, amino acid transport, iron uptake, and energy metabolism, although these genes did not form a clearly resolved STRING cluster (**Figure S2A**), whereas downregulated functions overlapped with those observed in strain SEM1.4Evo. In particular, genes encoding T6SS, various two-component sensory systems, carbohydrate-storage functions, glycosyl transferases, and fatty acid metabolism were consistently repressed in both adapted backgrounds (**Figure S2B**). This overlap indicates that the *gacA***^E197K^** and *fabB***^L77P^** mutations were sufficient to reproduce a large fraction of the evolved transcriptional program.

Direct comparison between strain SEM1.4 GacA^E197K^ FabB^L77P^ and strain SEM1.4Evo revealed a smaller set of differentially expressed genes, consistent with phenotypic convergence between the two backgrounds (**Figure 5C** and **5D**). Genes upregulated in the double mutant relative to strain SEM1.4Evo included the tricarboxylate transporter genes *tctABC*, the outer membrane porin *opdH*, and the acetate permease *actP-I*. These changes suggest that the double mutant may retain higher expression of transport functions linked to organic-acid uptake. Conversely, genes with lower expression in the double mutant included *scpC*, heat-shock and redox-associated genes, arginine catabolism genes, and nitrogen-assimilation functions. Overall, the transcriptomic data support a model in which acetate tolerance arises from *gacA*- and *fabB*-linked regulatory rewiring that reduces investment in costly stress-, secretion-, storage-, and envelope-associated programs. This transcriptional economy provides a plausible route to the higher biomass yield observed in the evolved and reverse-engineered strains.

### 3.7 Proteomic remodeling supports reduced stress burden and resource reallocation during acetate adaptation

Proteome-wide changes were quantified in the parental strain SEM1.4, strain SEM1.4Evo, and the reverse-engineered double mutant strain SEM1.4 GacA^E197K^ FabB^L77P^ during exponential growth in MSM with 100 mM acetate. Protein-abundance changes were analyzed as log_2_(fold-change) relative to the parental strain SEM1.4 or strain SEM1.4Evo, and differentially abundant proteins were visualized as volcano plots (**Figure 6A-C**). In agreement with the transcriptome, both acetate-adapted backgrounds showed extensive proteome remodeling relative to the parental strain SEM1.4 (**Figure S3A-B**), whereas direct comparison between the double mutant and strain SEM1.4Evo revealed only a limited set of differences (**Figure 6D**). This pattern further supports the notion that the two reverse-engineered mutations captured much of the evolved systems-level response.

**Figure 6.**
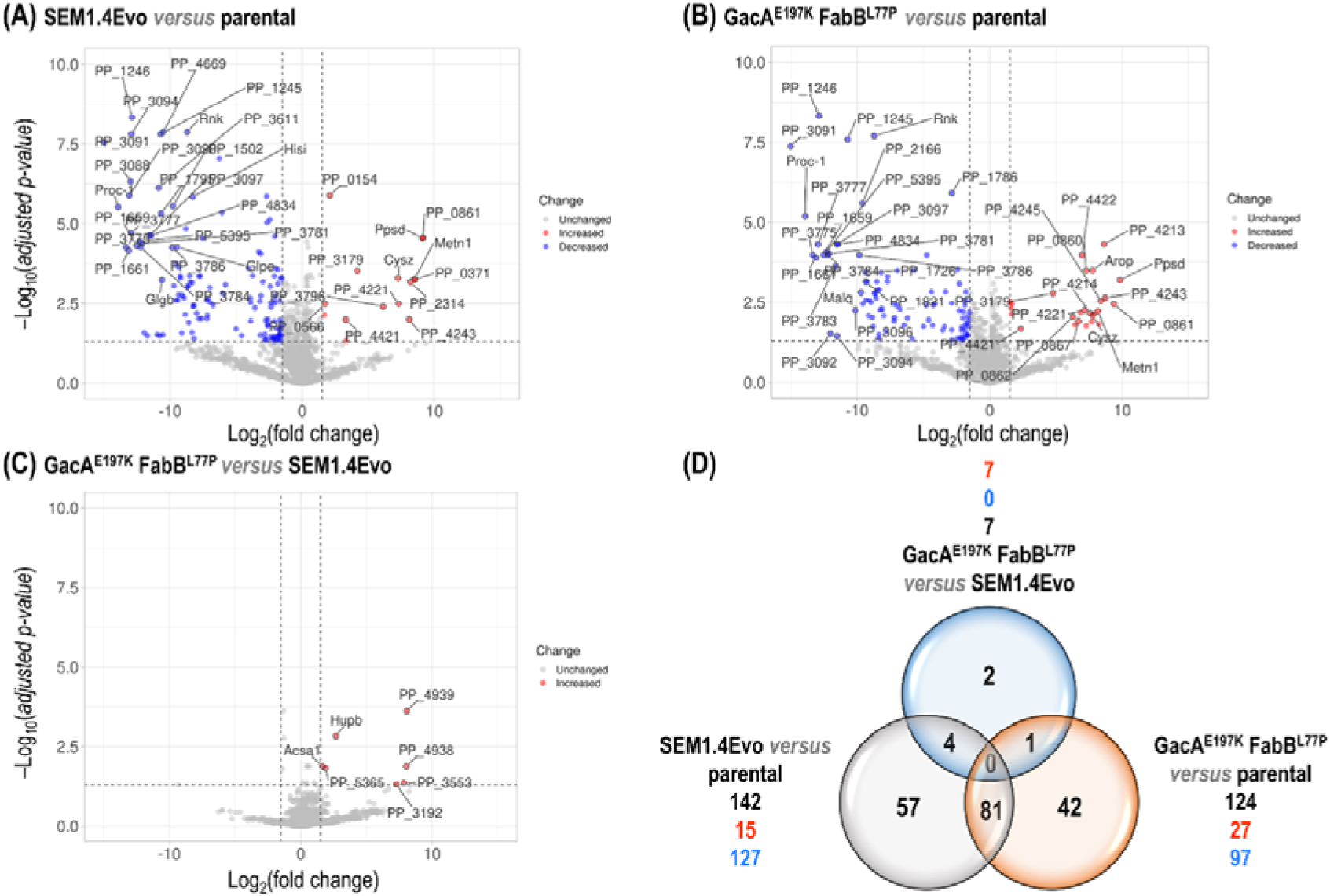
Proteomic analysis. Volcano plots showing differentially abundant proteins in *P. putida* SEM1.4Evo versus the parental strain (SEM1.4) **(A)**, reverse-engineered GacA^E197K^ FabB^L77P^ versus the parental strain **(B)**, and reverse-engineered GacA^E197K^ FabB^L77P^ versus SEM1.4Evo **(C)**. Four biological replicates were grown in MSM supplemented with 100 mM acetate and harvested during mid-exponential phase. Horizontal thresholds were set at 1.3 with p < 0.05, and vertical thresholds at −1.5 and +1.5 log_2_(fold change). **(D)** Venn diagram showing significantly regulated proteins shared among the comparisons in **(A–C)**. Red and blue numbers indicate upregulated and downregulated proteins, respectively, whereas dark gray indicates the total number of significantly regulated proteins in each comparison.

Several proteomic changes pointed to altered sulfur and methionine metabolism under acetate stress. MetN1, a methionine transport ATP-binding protein, and CysZ, a sulfate permease, were strongly increased in abundance in the adapted backgrounds. In contrast, proteins linked to organosulfur utilization and homocysteine metabolism, including SsuD, SsuF, GlpE, and MetY, were reduced. Methionine metabolism has been associated with protection against acetate-induced stress in bacteria, in part through its connection to homocysteine toxicity and sulfur assimilation. The protein-abundance pattern observed here suggests that acetate adaptation reshaped sulfur allocation and methionine-associated metabolism, potentially reducing the cost of stress management while preserving essential biosynthetic capacity.

A second major proteomic signature involved oxidative stress management. Proteins involved in reactive oxygen species detoxification, e.g., glutathione *S*-transferases, thiol peroxidase, catalase-peroxidase, superoxide dismutase, and cytochrome *c*-associated functions, were less abundant in strain SEM1.4Evo and in the double mutant than in the parental strain. This reduction is consistent with a lower oxidative stress burden in the adapted strains. The result aligns with the physiological data: the parental strain consumed acetate faster but converted it less efficiently into biomass, whereas evolved and reverse-engineered strains achieved higher biomass yields with lower acetate uptake. Reduced investment in oxidative-stress enzymes therefore supports the interpretation that less acetate-derived carbon and energy were spent on nonproductive stress compensation.

Proteomic changes also indicated remodeling of iron acquisition and fatty acid metabolism. Proteins associated with iron homeostasis and pyoverdine-linked functions were strongly and positively affected in the evolved and double mutant strains, consistent with the known connection between iron availability, oxidative stress, and the Gac regulatory network (89). In parallel, proteins involved in β-oxidation and fatty acid metabolism, including FadE, several acyl-CoA dehydrogenases, and cyclopropane-fatty-acyl-phospholipid synthase, were reduced in the adapted backgrounds. This pattern suggests that acetate adaptation decreased reliance on fatty acid turnover and membrane remodeling while redirecting acetyl-CoA and energetic resources toward growth-supporting metabolism. The recurrence of the *fabB***^L77P^** mutation further supports a role for lipid metabolism and membrane physiology in acetate tolerance.

Together, the proteomic data reinforce the resource-reallocation model suggested by the transcriptome. The evolved and reverse-engineered strains showed reduced abundance of proteins linked to energetically expensive stress responses, secretion programs, lipid turnover, and oxidative-stress defense (**Figure S4A-B**). These changes converge with the transcriptomic repression of T6SS, carbohydrate-storage genes, and fatty acid biosynthesis, and they help explain why acetate tolerance was associated with higher biomass yield rather than increased acetate uptake. In this model, the *gacA***^E197K^** and *fabB***^L77P^** mutations reprogram cellular resource allocation, reducing carbon and energy loss through stress-associated functions and improving biomass formation per acetate consumed.

### 3.8 ^13^C-metabolic flux analysis reveals carbon-conserving rewiring of acetate metabolism

The transcriptomic and proteomic profiles suggested that acetate adaptation reduced investment in energetically costly stress-response and non-essential cellular programs. To determine whether this regulatory shift was accompanied by changes in intracellular carbon distribution, we performed network-wide ^13^C-metabolic flux analysis during growth on 100 mM acetate. The parental strain SEM1.4, strain SEM1.4Evo, and the reverse-engineered double mutant strain SEM1.4 GacA^E197K^ FabB^L77P^ were cultivated with parallel acetate tracers, and labeling patterns from proteinogenic amino acids and cellular sugars were used to estimate central carbon fluxes (**Figure 7**).

**Figure 7.**
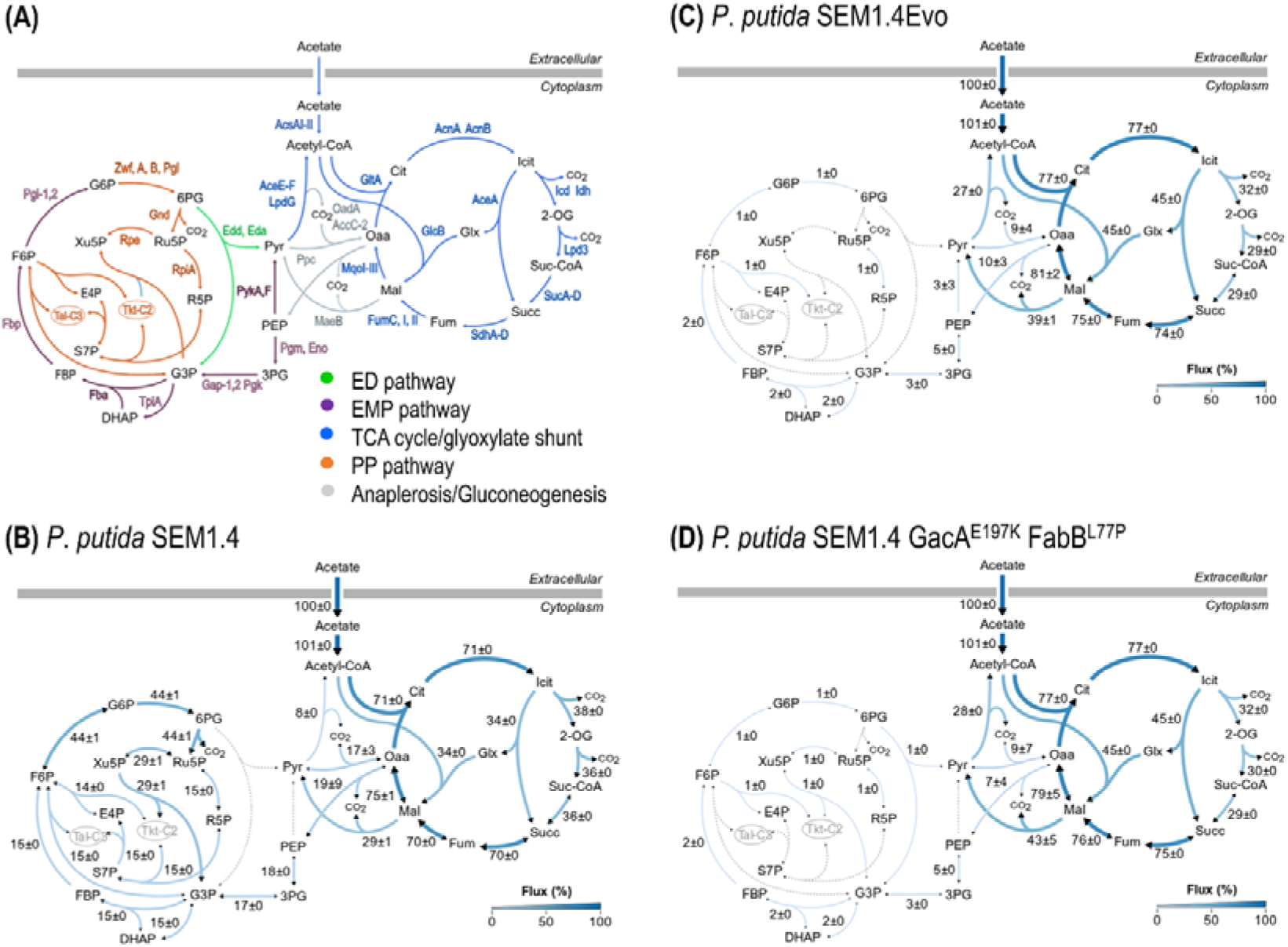
Fluxomic analysis. Acetate metabolism and key associated enzymes in P. putida KT2440 **(A)**. In vivo carbon flux distributions through central carbon metabolism in MSM cultures with 100 mM acetate in P. putida SEM1.4 **(B)**, SEM1.4Evo **(C)**, and the reverse-engineered GacA^E197K^ FabB^L77P^ strain **(D)**. Fluxes are expressed as percentages of the specific rates of acetate uptake (see Figure 4C). The tracer experiments were independently repeated three times; uncertainties associated with each flux indicate 95% confidence intervals. Fluxes are represented using a blue scale, with higher fluxes shown in darker blue and no flux indicated by gray dashed lines.

Flux distributions differed markedly between the parental strain SEM1.4 and the two acetate-adapted backgrounds. The parental strain showed substantial flux through the EDEMP cycle and pentose phosphate pathway, consistent with activation of cyclic central-carbon routes under acetate stress. Such cycling can support redox balancing and precursor supply (66), but it also increases carbon and energy expenditure under stress conditions. In contrast, strain SEM1.4Evo and the double mutant showed strongly reduced flux through these routes, indicating that acetate adaptation attenuated nonproductive central-metabolic cycling.

The adapted strains also redirected a larger fraction of acetate-derived carbon through carbon-conserving routes. During growth on acetate, ca. 50% of the carbon reaching the acetyl-CoA node entered the glyoxylate shunt in the parental strain SEM1.4 relative to the citrate/isocitrate branch, whereas this fraction increased to ca. 60% in strain SEM1.4Evo and the double mutant (**Figure 7**). Increased glyoxylate shunt flux is consistent with improved carbon retention because this route bypasses the CO_2_-generating steps of the tricarboxylic acid cycle and supports formation of C_4_ precursors required for gluconeogenesis and biomass synthesis (31). The evolved and reverse-engineered strains also showed increased pyruvate-shunt flux relative to the parental strain, suggesting an additional route to balance redox metabolism while maintaining growth-supporting carbon flow (101,102).

These fluxomic results provide an independent mechanistic explanation for the higher biomass yields observed in strain SEM1.4Evo and the double mutant. The parental strain consumed acetate faster but routed more carbon through stress-associated and cyclic central-metabolic activity, whereas the adapted strains consumed acetate more efficiently and retained more carbon in biomass. Together with transcriptomic repression of T6SS and carbohydrate-storage genes, and proteomic reduction of oxidative-stress-associated enzymes, the ^13^C-MFA data support a model in which the *gacA***^E197K^** and *fabB***^L77P^** mutations reduce costly stress-associated expenditure and redirect acetate carbon toward biomass formation. Acetate tolerance in the evolved background therefore reflects more efficient carbon use, rather than simply increased acetate uptake.

### 3.9 Structural modeling suggests that the gacA^E197K^ mutation perturbs the GacA DNA-binding domain

Drawing on the transcriptomic and proteomic observations, we further investigated the potential structural consequences of the *gacA***^E197K^** substitution. AlphaFold-based structural modeling indicated that residue 197 is located within the *C*-terminal LuxR/FixJ-type helix-turn-helix domain of GacA (**Figure 8A** and **8B**), a region associated with DNA recognition and transcriptional regulation (103,104). In the parental GacA model, Glu197 is positioned close to Thr193, with a predicted separation of ca. 2.9 to 3.1 Å (**Figure 8B**). This arrangement suggests a potential stabilizing interaction within the local structure of the helix-turn-helix domain.

**Figure 8.**
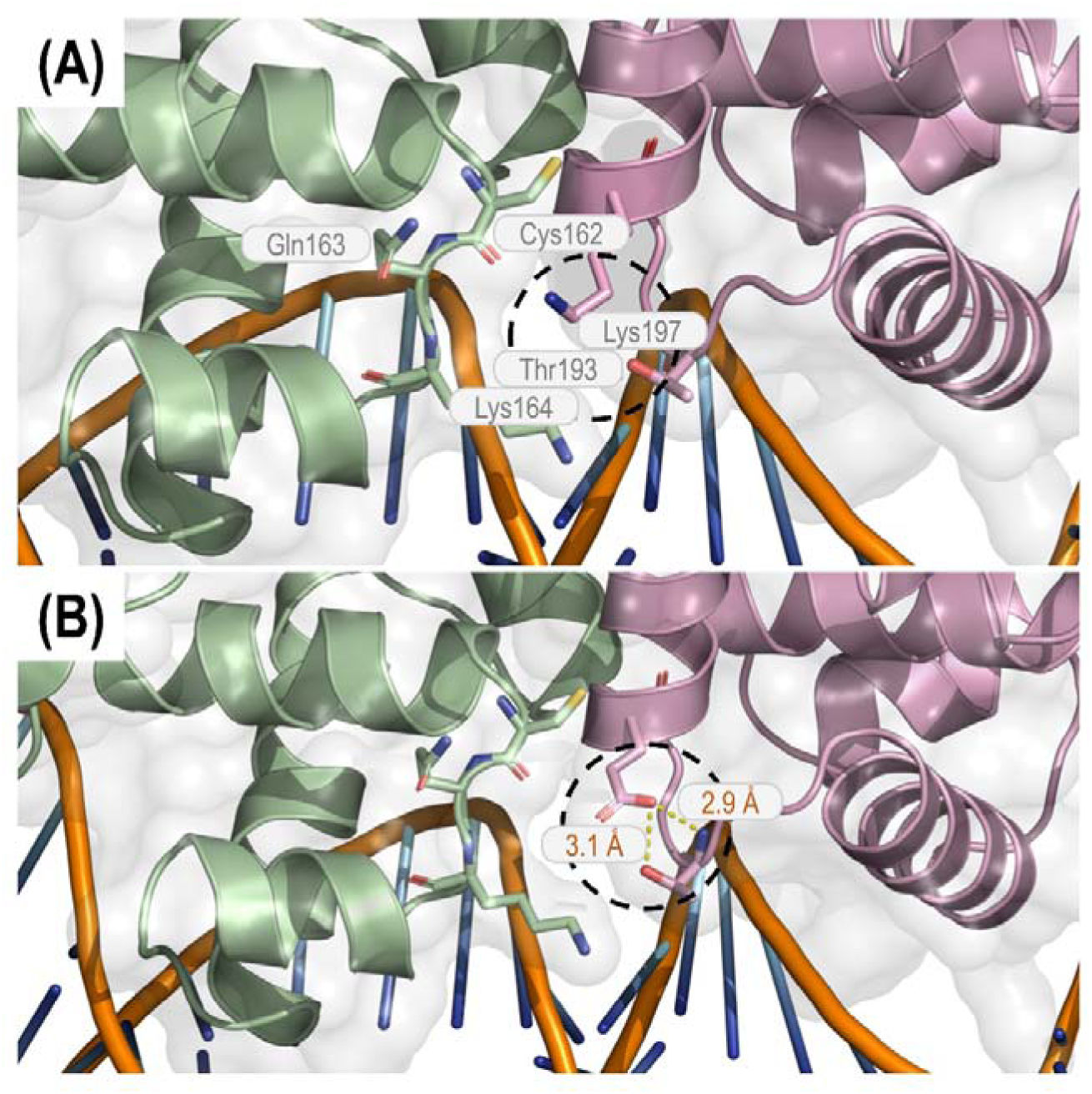
*In silico* structural analysis of the GacA protein of *P. putida*. Structural models of the GacA^E197K^ mutant **(A)** and wild-type GacA **(B)**, centered on residue 197 and the surrounding amino acids. Key residues (i.e., Cys162, Gln163, Lys164, Thr193, and Glu197/Lys197) are highlighted in the diagram. The residue selected for mutational analysis is indicated by a black dashed circle. Structures were visualized using PyMOL.

The E197K substitution introduces a charge reversal at this position and alters the orientation of the side chain in the mutant model (**Figure 8A**). No equivalent interaction line is predicted for Lys197 in the mutant structure, suggesting that the substitution perturbs the local electrostatic environment around residue 197. Although this modeling does not directly demonstrate altered DNA binding or protein-protein interactions, the predicted change in the local structural environment suggests that the substitution may impair GacA function. This interpretation is consistent with experimental evidence showing that inactivation of *gacS* or *gacA* in *P*. *putida* and *Pseudomonas stutzeri* can result in substantial fitness improvements (90). The extensive transcriptomic and proteomic changes observed in both strain SEM1.4Evo and the reverse-engineered double mutant are also compatible with reduced GacA-dependent regulation.

Together, these observations support the hypothesis that the *gacA***^E197K^** substitution contributes to the acetate-tolerant phenotype by modulating GacA function. In the context of the physiological and multi-omic data, attenuation of GacA-dependent regulation provides a plausible route to repress costly cellular programs, reduce stress-associated expenditure, and improve acetate-carbon conversion into biomass.

## 4. CONCLUSION

Acetate is an attractive C_2_ substrate for microbial bioproduction (4), but its use is constrained by toxicity, redox imbalance, the ATP cost of acetate activation, and poor carbon-use efficiency at elevated concentrations (12,105). This study used ALE to improve acetate tolerance in a genome-reduced strain of *P*. *putida* and then combined reverse engineering with transcriptomics, proteomics, and ^13^C-MFA to identify the systems-level basis of the evolved phenotype. The central outcome of this combined analysis is that acetate tolerance was not explained by faster acetate uptake. Instead, the evolved and reverse-engineered strains converted acetate into biomass more efficiently, indicating that adaptation improved carbon retention and reduced non-productive expenditure under acetate stress.

The evolved phenotype was largely associated with two recurrent mutations, *gacA***^E197K^** and *fabB***^L77P^**. Reverse engineering of these mutations into the parental strain SEM1.4 recapitulated most of the acetate-tolerant phenotype, including shorter lag phase, improved specific growth rate, and higher biomass yield than the parental strain. This result links acetate adaptation to two major cellular processes: global regulation through the GacS/GacA system and membrane-associated fatty acid metabolism through FabB. The GacS/GacA system controls broad adaptive programs in *Pseudomonas*, including secretion, motility, stress responses, and secondary metabolism (88,89). Attenuation of this system therefore provides a plausible route to reduce costly cellular functions during growth on acetate, as observed during evolution in the presence of aromatic compounds (87,95,96). The *fabB* mutation suggests that membrane physiology also contributes to acetate tolerance, possibly by modifying lipid metabolism, envelope properties, or the energetic cost of maintaining membrane integrity under weak-acid stress (92,93). A complementary study by Filbig et al. (91) examined acetate metabolism in wild-type *P*. *putida* KT2440 using a combination of rational engineering, tolerance-ALE, reverse engineering, transcriptomics, and 3-(3-hydroxyalkanoyloxy) alkanoic acid production as an acetyl-CoA–derived output. Despite differences in strain background and experimental scope, both studies converged on the GacS/GacA regulatory axis as a recurrent target for improving acetate performance. Furthermore, *crc* and *fleQ* were identified as additional adaptive nodes, and acetate-evolved strains retained useful production traits.

The interpretations emerging from the mutational landscape in our study are consistent with the “fear *versus* greed” framework described for bacterial adaptive evolution, in which growth-selected populations reduce investment in general stress and environmental defense programs while reallocating transcriptional and proteomic resources toward proliferation (106–108). Although this concept was formalized mainly through *E*. *coli* analyses, the acetate-evolved *P*. *putida* strains described here show an analogous systems-level shift. Repression of T6SS, carbohydrate storage functions, oxidative stress proteins, and lipid turnover pathways indicates a coordinated shift from a defensive, resource-intensive state toward a growth-efficient metabolic configuration. In this context, acetate evolution selected not simply for faster substrate consumption, but for a more “greedy” allocation strategy that increased biomass formation per unit of acetate consumed.

An informative physiological trait was the increase in biomass yield. The parental strain SEM1.4 consumed acetate faster than the evolved and double mutant strains, but this higher uptake rate did not translate into higher biomass formation. In contrast, strain SEM1.4Evo and the reverse-engineered double mutant showed lower acetate uptake rates but higher biomass yields. This pattern indicates that the parental strain dissipated a larger fraction of acetate-derived carbon and energy through stress responses, CO_2_-generating metabolism, maintenance demands, or futile central-metabolic cycling. The adapted strains instead retained a larger fraction of consumed acetate in biomass, and this phenotypic trait provides a mechanistic anchor for interpreting the omic datasets.

Transcriptomics supported this model by revealing strong repression of energetically expensive and non-essential cellular programs. T6SS genes, two-component sensory systems, carbohydrate storage functions, glycosyl transferases, and fatty acid biosynthesis genes were consistently downregulated in strain SEM1.4Evo and in the reverse-engineered double mutant. Repression of T6SS is particularly relevant because this secretion machinery imposes a high biosynthetic and energetic burden (109). Reduced expression of glycogen, trehalose, and maltose-associated genes further indicates decreased investment in storage and reserve-carbohydrate metabolism. Together, these changes suggest that acetate adaptation redirected transcriptional activity away from functions that do not directly support growth under acetate stress.

Proteomics provided an independent layer of support for resource reallocation. Proteins involved in oxidative-stress defense, including glutathione *S*-transferases, thiol peroxidase, catalase-peroxidase, superoxide dismutase, and cytochrome *c*-associated functions, were less abundant in the adapted strains than in the parental strain SEM1.4. Lower abundance of these proteins is consistent with a reduced oxidative stress burden or a decreased requirement for stress-compensatory metabolism. Proteomic changes in sulfur and methionine metabolism further suggest that acetate adaptation reshaped sulfur allocation and stress-protective metabolism. In parallel, reduced abundance of β-oxidation and fatty acid metabolism proteins points to altered lipid turnover and acetyl-CoA allocation (110). These proteomic signatures converge with the transcriptomic data and support the view that the adapted strains reduced expenditure on stress management and non-growth-associated functions.

The ^13^C-metabolic flux analysis data connected this regulatory and proteomic remodeling to central carbon metabolism. The parental strain SEM1.4 showed substantial flux through the EDEMP and pentose phosphate pathways during growth on acetate, consistent with activation of cyclic central-metabolic routes under stress (66). Such flux can support redox balancing and precursor supply, but it may also increase carbon and energy loss when maintained as a stress response. In contrast, strain SEM1.4Evo and the reverse-engineered double mutant showed strong attenuation of these routes and increased relative flux through the glyoxylate shunt. This shift is mechanistically important because the glyoxylate shunt bypasses CO_2_-generating steps of the TCA cycle and conserves carbon for biosynthesis (39). Increased glyoxylate-shunt flux therefore provides a direct explanation for improved biomass yield during growth on acetate.

The emerging model is that *gacA***^E197K^** and *fabB***^L77P^** reprogram acetate metabolism through coordinated regulatory, envelope, and metabolic effects. Attenuated GacA-dependent regulation reduces expression of costly secretion, sensing, and stress-associated programs. Changes associated with FabB likely affect membrane and fatty acid metabolism, reducing the burden of maintaining envelope homeostasis under weak-acid stress. Flux redistribution then channels acetate-derived carbon toward carbon-conserving metabolism, especially through the glyoxylate shunt, while reducing futile EDEMP cycling. These changes collectively improve biomass formation per acetate consumed.

This study also highlights the value of genome-reduced *P*. *putida* as a background for dissecting tolerance mechanisms. Genome reduction simplifies cellular architecture and can expose adaptation routes that improve growth and resource allocation under defined selective pressures. The acetate-tolerant derivatives generated here provide genetic targets for future strain engineering in acetate-based bioprocesses. In particular, modulation of GacA-dependent regulation, fatty acid metabolism, T6SS expression, and glyoxylate-shunt flux could be combined with product-forming pathways to increase carbon efficiency from acetate. Such strategies are relevant for biomanufacturing workflows that use acetate derived from biomass conversion, syngas fermentation, microbial electrosynthesis, or mixed-carbon waste streams (8).

Overall, acetate-driven evolution of *P*. *putida* SEM1.4 revealed a resource-efficient tolerance strategy centered on regulatory attenuation, reduced stress expenditure, and carbon-conserving flux redistribution. The evolved and reverse-engineered strains did not simply consume acetate faster, but they used acetate more efficiently. This distinction is critical for acetate-based bioproduction, where improved carbon retention and lower maintenance burden can be more valuable than increased substrate uptake alone.

## AUTHORS’ CONTRIBUTIONS

**Nicolás Gurdo:** conceptualization, methodology, investigation, formal analysis, validation, visualization, data curation, writing – original draft, writing – review and editing. **Aparajitha Srinivasan:** methodology, investigation, formal analysis, data curation, validation, writing – review and editing. **Tommaso Tagliani:** methodology, investigation, formal analysis. **Melanie Filbig:** investigation, methodology. **Nicolas T. Wirth:** methodology, software, formal analysis, validation, data curation. **Josefin Johnsen:** investigation, methodology. **Garret W. O’Connell:** investigation. **Stefano Donati:** formal analysis, methodology. **Enrico Orsi:** formal analysis, methodology, validation, data curation. **María V. G. Alván-Vargas:** investigation. **Yan Chen:** methodology. **Christopher J. Petzold:** methodology. **Matthew Blow:** methodology, resources, data curation, validation. **Thomas Eng:** resources, supervision, data curation, writing – review and editing. **Till Tiso:** methodology. **Lars M. Blank:** resources, supervision. **Adam Feist:** methodology, resources, supervision, funding acquisition. **Aindrila Mukhopadhyay:** conceptualization, resources, supervision, funding acquisition, project administration, writing – review and editing. **Pablo I. Nikel:** conceptualization, methodology, formal analysis, resources, supervision, funding acquisition, project administration, writing – review and editing.

## ACKNOWLEDGMENTS

The authors acknowledge the use of BioRender (https://biorender.com) for the creation of schematic diagrams in this manuscript. The authors initially wrote the complete manuscript and used ChatGPT for English grammar checking. The authors then reviewed and revised the text to ensure accuracy.

## DATA AVAILABILITY

All data used in this study are contained in the **Data S1** file. Raw RNA-Seq data are available at the JGI Genome Portal (https://genome.jgi.doe.gov/portal) under project ID 1445921. The mass spectrometry proteomics data have been deposited to the ProteomeXchange Consortium *via* the PRIDE partner repository (111) under dataset ID PXD075741. The DIA-NN software is available at https://github.com/vdemichev/DiaNN.

## REFERENCES

1. Schrader J, Schilling M, Holtmann D, Sell D, Filho MV, Marx A, Vorholt JA. 2009. Methanol-based industrial biotechnology: current status and future perspectives of methylotrophic bacteria. Trends Biotechnol 27:107–115. 10.1016/j.tibtech.2008.10.009.

2. Dürre P, Eikmanns BJ. 2015. C1-carbon sources for chemical and fuel production by microbial gas fermentation. Curr Opin Biotechnol 35:63–72.

3. Wendisch VF, Brito LF, Gil-López M, Hennig G, Pfeifenschneider J, Sgobba E, Veldmann KH. 2016. The flexible feedstock concept in Industrial Biotechnology: metabolic engineering of Escherichia coli, Corynebacterium glutamicum, Pseudomonas, Bacillus and yeast strains for access to alternative carbon sources. J Biotechnol 234:139–157. 10.1016/j.jbiotec.2016.07.022.

4. Kiefer D, Merkel M, Lilge L, Henkel M, Hausmann R. 2021. From acetate to bio-based products: underexploited potential for industrial biotechnology. Trends Biotechnol 39:397–411. 10.1016/j.tibtech.2020.09.004.

5. Novak K, Pflügl S. 2018. Towards biobased industry: acetate as a promising feedstock to enhance the potential of microbial cell factories. FEMS Microbiol Lett 365. 10.1093/femsle/fny226.

6. Zhang J, Liu Z, Wang Y, Yu B. 2024. Sustainable production of Lllhomoserine solely from CO_2_llderived acetate and formate by engineered E. coli strain. ACS Sust Chem Eng 12:18704–18711. 10.1021/acssuschemeng.4c08299.

7. Lim HG, Lee JH, Noh MH, Jung GY. 2018. Rediscovering acetate metabolism: its potential sources and utilization for biobased transformation into value-added chemicals. J Agric Food Chem 66:3998–4006. 10.1021/acs.jafc.8b00458.

8. Kim Y, Lama S, Agrawal D, Kumar V, Park S. 2021. Acetate as a potential feedstock for the production of value-added chemicals: metabolism and applications. Biotechnol Adv 49:107736. 10.1016/j.biotechadv.2021.107736.

9. Luli GW, Strohl WR. 1990. Comparison of growth, acetate production, and acetate inhibition of Escherichia coli strains in batch and fed-batch fermentations. Appl Environ Microbiol 56:1004–1011. 10.1128/aem.56.4.1004-1011.1990.

10. Kuang Z, Yan X, Yuan Y, Wang R, Zhu H, Wang Y, Li J, Ye J, Yue H, Yang X. 2024. Advances in stress-tolerance elements for microbial cell factories. Synth Syst Biotechnol 9:793–808. 10.1016/j.synbio.2024.06.008.

11. Lasko DR, Zamboni N, Sauer U. 2000. Bacterial response to acetate challenge: a comparison of tolerance among species. Appl Microbiol Biotechnol 54:243–247. 10.1007/s002530000339.

12. Trček J, Mira NP, Jarboe LR. 2015. Adaptation and tolerance of bacteria against acetic acid. Appl Microbiol Biotechnol 99:6215–6229. 10.1007/s00253-015-6762-3.

13. Pinhal S, Ropers D, Geiselmann J, de Jong H. 2019. Acetate metabolism and the inhibition of bacterial growth by acetate. J Bacteriol 201:e00147–19. 10.1128/jb.00147-19.

14. Chong H, Yeow J, Wang I, Song H, Jiang R. 2013. Improving acetate tolerance of Escherichia coli by rewiring its global regulator cAMP receptor protein (CRP). PLoS One 8:e77422. 10.1371/journal.pone.0077422.

15. Roe AJ, McLaggan D, Davidson I, O’Byrne C, Booth IR. 1998. Perturbation of anion balance during inhibition of growth of Escherichia coli by weak acids. J Bacteriol 180:767–772. 10.1128/jb.180.4.767-772.1998.

16. Nikel PI, de Lorenzo V. 2018. Pseudomonas putida as a functional chassis for industrial biocatalysis: From native biochemistry to trans-metabolism. Metab Eng 50:142–155. 10.1016/j.ymben.2018.05.005.

17. Volke DC, Calero P, Nikel PI. 2020. Pseudomonas putida. Trends Microbiol 28:512–513. 10.1016/j.tim.2020.02.015.

18. Weimer A, Kohlstedt M, Volke DC, Nikel PI, Wittmann C. 2020. Industrial biotechnology of Pseudomonas putida: advances and prospects. Appl Microbiol Biotechnol 104:7745–7766. 10.1007/s00253-020-10811-9.

19. de Lorenzo V, Pérez-Pantoja D, Nikel PI. 2024. Pseudomonas putida KT2440: the long journey of a soil-dweller to become a synthetic biology chassis. J Bacteriol 206:e00136–24. 10.1128/jb.00136-24.

20. Lieder S, Nikel PI, de Lorenzo V, Takors R. 2015. Genome reduction boosts heterologous gene expression in Pseudomonas putida. Microb Cell Fact 14:23. 10.1186/s12934-015-0207-7.

21. Volke DC, Friis L, Wirth NT, Turlin J, Nikel PI. 2020. Synthetic control of plasmid replication enables target- and self-curing of vectors and expedites genome engineering of Pseudomonas putida. Metab Eng Commun 10:e00126. 10.1016/j.mec.2020.e00126.

22. Wirth NT, Rohr K, Danchin A, Nikel PI. 2023. Recursive genome engineering decodes the evolutionary origin of an essential thymidylate kinase activity in Pseudomonas putida KT2440. mBio 14:e01081–23. 10.1128/mbio.01081-23.

23. Wynands B, Otto M, Runge N, Preckel S, Polen T, Blank LM, Wierckx N. 2019. Streamlined Pseudomonas taiwanensis VLB120 chassis strains with improved bioprocess features. ACS Synth Biol 8:2036–2050. 10.1021/acssynbio.9b00108.

24. Ma S, Su T, Lu X, Qi Q. 2024. Bacterial genome reduction for optimal chassis of synthetic biology: a review. Crit Rev Biotechnol 44:660–673. 10.1080/07388551.2023.2208285.

25. Calero P, Nikel PI. 2019. Chasing bacterial chassis for metabolic engineering: a perspective review from classical to non-traditional microorganisms. Microb Biotechnol 12:98–124. 10.1111/1751-7915.13292.

26. Yus E, Maier T, Michalodimitrakis K, van Noort V, Yamada T, Chen WH, Wodke JA, Güell M, Martínez S, Bourgeois R, Kühner S, Raineri E, Letunic I, Kalinina OV, Rode M, Herrmann R, Gutiérrez-Gallego R, Russell RB, Gavin AC, Bork P, Serrano L. 2009. Impact of genome reduction on bacterial metabolism and its regulation. Science 326:1263–1268. 10.1126/science.1177263.

27. Díaz-Guerra M, Esteban M, Martínez JL. 1997. Growth of Escherichia coli in acetate as a sole carbon source is inhibited by ankyrin-like repeats present in the 2’,5’-linked oligoadenylate-dependent human RNase L enzyme. FEMS Microbiol Lett 149:107–113. 10.1111/j.1574-6968.1997.tb10316.x.

28. Roe AJ, O’Byrne C, McLaggan D, Booth IR. 2002. Inhibition of Escherichia coli growth by acetic acid: a problem with methionine biosynthesis and homocysteine toxicity. Microbiology 148:2215–2222. 10.1099/00221287-148-7-2215.

29. Seong W, Han GH, Lim HS, Baek JI, Kim SJ, Kim D, Kim SK, Lee H, Kim H, Lee SG, Lee DH. 2020. Adaptive laboratory evolution of Escherichia coli lacking cellular byproduct formation for enhanced acetate utilization through compensatory ATP consumption. Metab Eng 62:249–259. 10.1016/j.ymben.2020.09.005.

30. Viana de Siqueira GM, Eng T, Mukhopadhyay A, Guazzaroni ME. 2025. Differences in GenBank and RefSeq annotations may affect genomics data interpretation for Pseudomonas putida KT2440. mSphere 10:e00391–25. 10.1128/msphere.00391-25.

31. Turlin J, Alván-Vargas MVG, Puiggené Ò, Donati S, Wenk S, Nikel PI. 2025. Synthetic C_1_ metabolism in Pseudomonas putida enables strict formatotrophy and methylotrophy via the reductive glycine pathway. mBio 16:01976–25. 10.1128/mbio.01976-25.

32. Puiggené Ò, Muñoz-Triviño J, Civil-Ferrer L, Gille L, Schulz-Mirbach H, Bergen D, Erb TJ, Ebert BE, Nikel PI. 2025. Systematic engineering of synthetic serine cycles in Pseudomonas putida uncovers emergent topologies for methanol assimilation. Trends Biotechnol 43:2539–2565. 10.1016/j.tibtech.2025.06.001.

33. Choe D, Lee JH, Yoo M, Hwang S, Sung BH, Cho S, Palsson BØ, Kim SC, Cho BK. 2019. Adaptive laboratory evolution of a genome-reduced Escherichia coli. Nat Commun 10:935. 10.1038/s41467-019-08888-6.

34. Dragosits M, Mattanovich D. 2013. Adaptive laboratory evolution – Principles and applications for biotechnology. Microb Cell Fact 12:64. 10.1186/1475-2859-12-64.

35. Sandberg TE, Salazar MJ, Weng LL, Palsson BØ, Feist AM. 2019. The emergence of adaptive laboratory evolution as an efficient tool for biological discovery and industrial biotechnology. Metab Eng 56:1–16. 10.1016/j.ymben.2019.08.004.

36. Fernández-Cabezón L, Cros A, Nikel PI. 2019. Evolutionary approaches for engineering industrially-relevant phenotypes in bacterial cell factories. Biotechnol J 14:1800439. 10.1002/biot.201800439.

37. Horinouchi T, Furusawa C. 2020. Understanding metabolic adaptation by using bacterial laboratory evolution and trans-omics analysis. Biophys Rev 12:677–682. 10.1007/s12551-020-00695-4.

38. Sandberg TE, Lloyd CJ, Palsson BØ, Feist AM. 2017. Laboratory evolution to alternating substrate environments yields distinct phenotypic and genetic adaptive strategies. Appl Environ Microbiol 83:e00410–17.

39. Turlin J, Dronsella B, De Maria A, Lindner SN, Nikel PI. 2022. Integrated rational and evolutionary engineering of genome-reduced Pseudomonas putida strains promotes synthetic formate assimilation. Metab Eng 74:191–205. 10.1016/j.ymben.2022.10.008.

40. Phaneuf PV, Gosting D, Palsson BØ, Feist AM. 2019. ALEdb 1.0: a database of mutations from adaptive laboratory evolution experimentation. Nucleic Acids Res 47:D1164–D1171. 10.1093/nar/gky983.

41. Deatherage DE, Barrick JE. 2014. Identification of mutations in laboratory-evolved microbes from next-generation sequencing data using breseq. Methods Mol Biol 1151:165–188. 10.1007/978-1-4939-0554-6_12.

42. Belda E, van Heck RGA, López-Sánchez MJ, Cruveiller S, Barbe V, Fraser C, Klenk HP, Petersen J, Morgat A, Nikel PI, Vallenet D, Rouy Z, Sekowska A, Martins dos Santos VAP, de Lorenzo V, Danchin A, Médigue C. 2016. The revisited genome of Pseudomonas putida KT2440 enlightens its value as a robust metabolic chassis. Environ Microbiol 18:3403–3424. 10.1111/1462-2920.13230.

43. LaCroix RA, Palsson BØ, Feist AM. 2017. A model for designing adaptive laboratory evolution experiments. Appl Environ Microbiol 83:e03115–16. 10.1128/AEM.03115-16.

44. Cavaleiro AM, Kim SH, Seppälä S, Nielsen MT, Nørholm MH. 2015. Accurate DNA assembly and genome engineering with optimized uracil excision cloning. ACS Synth Biol 4:1042–1046. 10.1021/acssynbio.5b00113.

45. Genee HJ, Bonde MT, Bagger FO, Jespersen JB, Sommer MOA, Wernersson R, Olsen LR. 2015. Software-supported USER cloning strategies for site-directed mutagenesis and DNA assembly. ACS Synth Biol 4:342–349. 10.1021/sb500194z.

46. Nikel PI, Pettinari MJ, Ramírez MC, Galvagno MA, Méndez BS. 2008. Escherichia coli arcA mutants: metabolic profile characterization of microaerobic cultures using glycerol as a carbon source. J Mol Microbiol Biotechnol 15:48–54. 10.1159/000111992.

47. Wirth NT, Kozaeva E, Nikel PI. 2020. Accelerated genome engineering of Pseudomonas putida by I-SceI―mediated recombination and CRISPR-Cas9 counterselection. Microb Biotechnol 13:233–249. 10.1111/1751-7915.13396.

48. Wirth NT, Funk J, Donati S, Nikel PI. 2023. QurvE: user-friendly software for the analysis of biological growth and fluorescence data. Nat Protoc 18:2401–2403. 10.1038/s41596-023-00850-7.

49. Federici F, Luppino F, Aguilar-Vilar C, Mazaraki ME, Petersen LB, Ahonen L, Nikel PI. 2025. CIFR (Clone-Integrate-Flip-out-Repeat): a toolset for iterative genome and pathway engineering of Gram-negative bacteria. Metab Eng 88:180–195. 10.1016/j.ymben.2025.01.001.

50. Kim D, Langmead B, Salzberg SL. 2015. HISAT: a fast spliced aligner with low memory requirements. Nat Methods 12:357–360. 10.1038/nmeth.3317.

51. Ramírez F, Dündar F, Diehl S, Grüning BA, Manke T. 2014. deepTools: a flexible platform for exploring deep-sequencing data. Nucleic Acids Res 42:W187–W191. 10.1093/nar/gku365.

52. Liao Y, Smyth GK, Shi W. 2014. featureCounts: an efficient general purpose program for assigning sequence reads to genomic features. Bioinformatics 30:923–930. 10.1093/bioinformatics/btt656.

53. Love MI, Huber W, Anders S. 2014. Moderated estimation of fold change and dispersion for RNA-seq data with DESeq2. Genome Biol 15:550. 10.1186/s13059-014-0550-8.

54. Turlin J, Puiggené Ò, Donati S, Wirth NT, Nikel PI. 2023. Core and auxiliary functions of one-carbon metabolism in Pseudomonas putida exposed by a systems-level analysis of transcriptional and physiological responses. mSystems 8:e00004–23. 10.1128/msystems.00004-23.

55. Kozaeva E, Volkova S, Matos MRA, Mezzina MP, Wulff T, Volke DC, Nielsen LK, Nikel PI. 2021. Model-guided dynamic control of essential metabolic nodes boosts acetyl-coenzyme A–dependent bioproduction in rewired Pseudomonas putida. Metab Eng 67:373–386. 10.1016/j.ymben.2021.07.014.

56. Silva JC, Denny R, Dorschel C, Gorenstein MV, Li GZ, Richardson K, Wall D, Geromanos SJ. 2006. Simultaneous qualitative and quantitative analysis of the Escherichia coli proteome: a SWEET tale. Mol Cell Proteomics 5:589–607. 10.1074/mcp.M500321-MCP200.

57. Ahrné E, Ohta Y, Nikitin F, Scherl A, Lisacek F, Müller M. 2011. An improved method for the construction of decoy peptide MS/MS spectra suitable for the accurate estimation of false discovery rates. Proteomics 11:4085–4095. 10.1002/pmic.201000665.

58. Gurdo N, Taylor Parkins SK, Fricano M, Wulff T, Nielsen LK, Nikel PI. 2023. Protocol for absolute quantification of proteins in Gram-negative bacteria based on QconCAT-based labeled peptides. STAR Protoc 4:102060. 10.1016/j.xpro.2023.102060.

59. Goedhart J, Luijsterburg MS. 2020. VolcaNoseR is a web app for creating, exploring, labeling and sharing volcano plots. Sci Rep 10:20560. 10.1038/s41598-020-76603-3.

60. Nikel PI, Zhu J, San KY, Méndez BS, Bennett GN. 2009. Metabolic flux analysis of Escherichia coli creB and arcA mutants reveals shared control of carbon catabolism under microaerobic growth conditions. J Bacteriol 191:5538–5548. 10.1128/JB.00174-09.

61. Zamboni N, Fendt SM, Rühl M, Sauer U. 2009. ^13^C-based metabolic flux analysis. Nat Protoc 4:878–92. 10.1038/nprot.2009.58.

62. Kiefer P, Heinzle E, Zelder O, Wittmann C. 2004. Comparative metabolic flux analysis of lysine-producing Corynebacterium glutamicum cultured on glucose or fructose. Appl Environ Microbiol 70:229–239. 10.1128/aem.70.1.229-239.2004.

63. Kutuzova S, Colaianni P, Röst H, Sachsenberg T, Alka O, Kohlbacher O, Burla B, Torta F, Schrübbers L, Kristensen M, Nielsen L, Herrgård MJ, McCloskey D. 2020. SmartPeak automates targeted and quantitative metabolomics data processing. Anal Chem 92:15968–15974. 10.1021/acs.analchem.0c03421.

64. Young JD. 2014. INCA: a computational platform for isotopically non-stationary metabolic flux analysis. Bioinformatics 30:1333–5. 10.1093/bioinformatics/btu015.

65. Nikel PI, Chavarría M, Fuhrer T, Sauer U, de Lorenzo V. 2015. Pseudomonas putida KT2440 strain metabolizes glucose through a cycle formed by enzymes of the Entner-Doudoroff, Embden-Meyerhof-Parnas, and pentose phosphate pathways. J Biol Chem 290:25920–25932. 10.1074/jbc.M115.687749.

66. Nikel PI, Fuhrer T, Chavarría M, Sánchez-Pascuala A, Sauer U, de Lorenzo V. 2021. Reconfiguration of metabolic fluxes in Pseudomonas putida as a response to sub-lethal oxidative stress. ISME J 15:1751–1766. 10.1038/s41396-020-00884-9.

67. Kohlstedt M, Wittmann C. 2019. GC-MS-based ^13^C metabolic flux analysis resolves the parallel and cyclic glucose metabolism of Pseudomonas putida KT2440 and Pseudomonas aeruginosa PAO1. Metab Eng 54:35–53. 10.1016/j.ymben.2019.01.008.

68. Nogales J, Mueller J, Gudmundsson S, Canalejo FJ, Duque E, Monk J, Feist AM, Ramos JL, Niu W, Palsson BØ. 2020. High-quality genome-scale metabolic modelling of Pseudomonas putida highlights its broad metabolic capabilities. Environ Microbiol 22:255–269. 10.1111/1462-2920.14843.

69. Bujdoš D, Popelářová B, Volke DC, Nikel PI, Sonnenschein N, Dvořák P. 2023. Engineering of Pseudomonas putida for accelerated co-utilization of glucose and cellobiose yields aerobic overproduction of pyruvate explained by an upgraded metabolic model. Metab Eng 75:29–46. 10.1016/j.ymben.2022.10.011.

70. Czajka JJ, Banerjee D, Eng T, Menasalvas J, Yan C, Muñoz NM, Poirier BC, Kim YM, Baker SE, Tang YJ, Mukhopadhyay A. 2022. Tuning a high performing multiplexed-CRISPRi Pseudomonas putida strain to further enhance indigoidine production. Metab Eng Commun 15:e00206. 10.1016/j.mec.2022.e00206.

71. Antoniewicz MR, Kelleher JK, Stephanopoulos G. 2006. Determination of confidence intervals of metabolic fluxes estimated from stable isotope measurements. Metab Eng 8:324–337. 10.1016/j.ymben.2006.01.004.

72. Szklarczyk D, Kirsch R, Koutrouli M, Nastou K, Mehryary F, Hachilif R, Gable AL, Fang T, Doncheva NT, Pyysalo S, Bork P, Jensen LJ, von Mering C. 2023. The STRING database in 2023: protein-protein association networks and functional enrichment analyses for any sequenced genome of interest. Nucleic Acids Res 51:D638–D646. 10.1093/nar/gkac1000.

73. Martino RA, Volke DC, Tenaglia AH, Tribelli PM, Nikel PI, Smania AM. 2025. Genetic dissection of cyclic di-GMP signalling in Pseudomonas aeruginosa via systematic diguanylate cyclase disruption. Microb Biotechnol 18:e70137. 10.1111/1751-7915.70137.

74. Jumper J, Evans R, Pritzel A, Green T, Figurnov M, Ronneberger O, Tunyasuvunakool K, Bates R, Žídek A, Potapenko A, Bridgland A, Meyer C, Kohl SAA, Ballard AJ, Cowie A, Romera-Paredes B, Nikolov S, Jain R, Adler J, Back T, Petersen S, Reiman D, Clancy E, Zielinski M, Steinegger M, Pacholska M, Berghammer T, Bodenstein S, Silver D, Vinyals O, Senior AW, Kavukcuoglu K, Kohli P, Hassabis D. 2021. Highly accurate protein structure prediction with AlphaFold. Nature 596:583–589. 10.1038/s41586-021-03819-2.

75. Aguirre-Plans J, Meseguer A, Molina-Fernandez R, Marín-López MA, Jumde G, Casanova K, Bonet J, Fornes O, Fernandez-Fuentes N, Oliva B. 2021. SPServer: split-statistical potentials for the analysis of protein structures and protein-protein interactions. BMC Bioinformatics 22:4. 10.1186/s12859-020-03770-5.

76. Wiederstein M, Sippl MJ. 2007. ProSA-web: interactive web service for the recognition of errors in three-dimensional structures of proteins. Nucleic Acids Res 35:W407–W410. 10.1093/nar/gkm290.

77. Winsor GL, Griffiths EJ, Lo R, Dhillon BK, Shay JA, Brinkman FS. 2016. Enhanced annotations and features for comparing thousands of Pseudomonas genomes in the Pseudomonas genome database. Nucleic Acids Res 44:D646–D653. 10.1093/nar/gkv1227.

78. Paysan-Lafosse T, Blum M, Chuguransky S, Grego T, Pinto BL, Salazar Gustavo A, Bileschi Maxwell L, Bork P, Bridge A, Colwell L, Gough J, Haft Daniel H, Letunić I, Marchler-Bauer A, Mi H, Natale Darren A, Orengo Christine A, Pandurangan Arun P, Rivoire C, Sigrist CJA, Sillitoe I, Thanki N, Thomas PD, Tosatto SCE, Wu Cathy H, Bateman A. 2023. InterPro in 2022. Nucleic Acids Res 51:D418–D427. 10.1093/nar/gkac993.

79. Waterhouse A, Bertoni M, Bienert S, Studer G, Tauriello G, Gumienny R, Heer FT, de Beer TAP, Rempfer C, Bordoli L, Lepore R, Schwede T. 2018. SWISS-MODEL: Homology modelling of protein structures and complexes. Nucleic Acids Res 46:W296–W303. 10.1093/nar/gky427.

80. Kirkpatrick C, Maurer LM, Oyelakin NE, Yoncheva YN, Maurer R, Slonczewski JL. 2001. Acetate and formate stress: opposite responses in the proteome of Escherichia coli. J Bacteriol 183:6466–6477. 10.1128/JB.183.21.6466-6477.2001.

81. Guan N, Liu L. 2020. Microbial response to acid stress: mechanisms and applications. Appl Microbiol Biotechnol 104:51–65. 10.1007/s00253-019-10226-1.

82. Rajaraman E, Agrawal A, Crigler J, Seipelt-Thiemann R, Altman E, Eiteman MA. 2016. Transcriptional analysis and adaptive evolution of Escherichia coli strains growing on acetate. Appl Microbiol Biotechnol 100:7777–7785. 10.1007/s00253-016-7724-0.

83. Fernández-Sandoval MT, Huerta-Beristain G, Trujillo-Martinez B, Bustos P, González V, Bolívar F, Gosset G, Martinez A. 2012. Laboratory metabolic evolution improves acetate tolerance and growth on acetate of ethanologenic Escherichia coli under non-aerated conditions in glucose-mineral medium. Appl Microbiol Biotechnol 96:1291–1300. 10.1007/s00253-012-4177-y.

84. Hua S, Wang Y, Wang L, Zhou Q, Li Z, Liu P, Wang K, Zhu Y, Han D, Yu Y. 2024. Regulatory mechanisms of acetic acid, ethanol and high temperature tolerances of acetic acid bacteria during vinegar production. Microb Cell Fact 23:324. 10.1186/s12934-024-02602-y.

85. Ackermann YS, de Witt J, Mezzina MP, Schroth C, Polen T, Nikel PI, Wynands B, Wierckx N. 2024. Bio-upcycling of even and uneven medium-chain-length diols and dicarboxylates to polyhydroxyalkanoates using engineered Pseudomonas putida. Microb Cell Fact 23:54. 10.1186/s12934-024-02310-7.

86. Martínez-García E, Nikel PI, Aparicio T, de Lorenzo V. 2014. Pseudomonas 2.0: genetic upgrading of P. putida KT2440 as an enhanced host for heterologous gene expression. Microb Cell Fact 13:159. 10.1186/s12934-014-0159-3.

87. Mohamed ET, Werner AZ, Salvachúa D, Singer CA, Szostkiewicz K, Jiménez-Díaz RM, Eng T, Radi MS, Simmons BA, Mukhopadhyay A, Herrgård MJ, Singer SW, Beckham GT, Feist AM. 2020. Adaptive laboratory evolution of Pseudomonas putida KT2440 improves p-coumaric and ferulic acid catabolism and tolerance. Metab Eng Commun 11:e00143. 10.1016/j.mec.2020.e00143.

88. Hassan KA, Johnson A, Shaffer BT, Ren Q, Kidarsa TA, Elbourne LD, Hartney S, Duboy R, Goebel NC, Zabriskie TM, Paulsen IT, Loper JE. 2010. Inactivation of the GacA response regulator in Pseudomonas fluorescens Pf-5 has far-reaching transcriptomic consequences. Environ Microbiol 12:899–915. 10.1111/j.1462-2920.2009.02134.x.

89. Song H, Li Y, Wang Y. 2023. Two-component system GacS/GacA, a global response regulator of bacterial physiological behaviors. Eng Microbiol 3:100051. 10.1016/j.engmic.2022.100051.

90. Eng T, Banerjee D, Lau AK, Bowden E, Herbert RA, Trinh J, Prahl JP, Deutschbauer A, Tanjore D, Mukhopadhyay A. 2021. Engineering Pseudomonas putida for efficient aromatic conversion to bioproduct using high throughput screening in a bioreactor. Metab Eng 66:229–238. 10.1016/j.ymben.2021.04.015.

91. Filbig M, Wachtendonk L, Hampe L, Bator I, Johnsen J, Mohamed ET, Gurdo N, Parschau J, Nikel PI, Feist AM, Tiso T, Blank LM. 2026. Improving acetate metabolism of P. putida KT2440 by evolutionary and rational engineering. bioRxiv, DOI: 10.64898/2026.08.21.746131.

92. Hoang TT, Schweizer HP. 1997. Fatty acid biosynthesis in Pseudomonas aeruginosa: cloning and characterization of the fabAB operon encoding β-hydroxyacyl-acyl carrier protein dehydratase (FabA) and β-ketoacyl-acyl carrier protein synthase I (FabB). J Bacteriol 179:5326–5332. 10.1128/jb.179.17.5326-5332.1997.

93. Yadrykhins’ky V, Georgiou C, Brenk R. 2021. Crystal structure of Pseudomonas aeruginosa FabB C161A, a template for structure-based design for new antibiotics. F1000Res 10:74018. 10.12688/f1000research.74018.2.

94. Kim S, Ha J, Lee H, Lee S, Lee J, Choi Y, Oh H, Yoon Y, Choi KH. 2019. Role of Pseudomonas aeruginosa DesB in adaptation to osmotic stress. J Food Prot 82:1278–1282. 10.4315/0362-028x.Jfp-18-507.

95. Bleem AC, Kuatsjah E, Johnsen J, Mohamed ET, Alexander WG, Kellermyer ZA, Carroll AL, Rossi R, Schlander IB, Peabody V GL, Guss AM, Feist AM, Beckham GT. 2024. Evolution and engineering of pathways for aromatic O-demethylation in Pseudomonas putida KT2440. Metab Eng 84:145–157. 10.1016/j.ymben.2024.06.009.

96. Werner AZ, Avina YC, Johnsen J, Bratti F, Alt HM, Mohamed ET, Clare R, Mand TD, Guss AM, Feist AM, Beckham GT. 2025. Adaptive laboratory evolution and genetic engineering improved terephthalate utilization in Pseudomonas putida KT2440. Metab Eng 88:196–205. 10.1016/j.ymben.2024.12.006.

97. Bull CT, Duffy B, Voisard C, Défago G, Keel C, Haas D. 2001. Characterization of spontaneous gacS and gacA regulatory mutants of Pseudomonas fluorescens biocontrol strain CHA0. Antonie van Leeuwenhoek 79:327–336. 10.1023/A:1012061014717.

98. Duffy BK, Défago G. 2000. Controlling instability in gacS-gacA regulatory genes during inoculant production of Pseudomonas fluorescens biocontrol strains. Appl Environ Microbiol 66:3142–3150. 10.1128/AEM.66.8.3142-3150.2000.

99. van den Broek D, Chin-A-Woeng TFC, Bloemberg GV, Lugtenberg BJJ. 2005. Molecular nature of spontaneous modifications in gacS which cause colony phase variation in Pseudomonas sp. strain PCL1171. J Bacteriol 187:593–600. 10.1128/jb.187.2.593-600.2005.

100. Yan Q, Lopes LD, Shaffer BT, Kidarsa TA, Vining O, Philmus B, Song C, Stockwell VO, Raaijmakers JM, McPhail KL, Andreote FD, Chang JH, Loper JE. 2018. Secondary metabolism and interspecific competition affect accumulation of spontaneous mutants in the GacS-GacA regulatory system in Pseudomonas protegens. mBio 9:e01845–17. 10.1128/mbio.01845-17.

101. Chavarría M, Goñi-Moreno A, de Lorenzo V, Nikel PI. 2016. A metabolic widget adjusts the phosphoenolpyruvate-dependent fructose influx in Pseudomonas putida. mSystems 1:e00154–16. 10.1128/mSystems.00154-16.

102. Chavarría M, Kleijn RJ, Sauer U, Pflüger-Grau K, de Lorenzo V. 2012. Regulatory tasks of the phosphoenolpyruvate-phosphotransferase system of Pseudomonas putida in central carbon metabolism. mBio 3:e00028–12. 10.1128/mBio.00028-12.

103. Kahn D, Ditta G. 1991. Modular structure of FixJ: homology of the transcriptional activator domain with the –35 binding domain of σ factors. Mol Microbiol 5:987–997. 10.1111/j.1365-2958.1991.tb00774.x.

104. Sitnikov DM, Schineller JB, Baldwin TO. 1995. Transcriptional regulation of bioluminesence genes from Vibrio fischeri. Mol Microbiol 17:801–812. 10.1111/j.1365-2958.1995.mmi_17050801.x.

105. Wolfe AJ. 2005. The acetate switch. Microbiol Mol Biol Rev 69:12–50. 10.1128/mmbr.69.1.12-50.2005.

106. Utrilla J, O’Brien Edward J, Chen K, McCloskey D, Cheung J, Wang H, Armenta-Medina D, Feist Adam M, Palsson BØ. 2016. Global rebalancing of cellular resources by pleiotropic point mutations illustrates a multi-scale mechanism of adaptive evolution. Cell Syst 2:260–271. 10.1016/j.cels.2016.04.003.

107. Tan J, Sastry AV, Fremming KS, Bjørn SP, Hoffmeyer A, Seo S, Voldborg BG, Palsson BØ. 2020. Independent component analysis of E. coli’s transcriptome reveals the cellular processes that respond to heterologous gene expression. Metab Eng 61:360–368. 10.1016/j.ymben.2020.07.002.

108. Dalldorf C, Rychel K, Szubin R, Hefner Y, Patel A, Zielinski DC, Palsson BØ. 2024. The hallmarks of a tradeoff in transcriptomes that balances stress and growth functions. mSystems 9:e00305–24. 10.1128/msystems.00305-24.

109. Bernal P, Allsopp LP, Filloux A, Llamas MA. 2017. The Pseudomonas putida T6SS is a plant warden against phytopathogens. ISME J 11:972–987. 10.1038/ismej.2016.169.

110. Fujita Y, Matsuoka H, Hirooka K. 2007. Regulation of fatty acid metabolism in bacteria. Mol Microbiol 66:829–39. 10.1111/j.1365-2958.2007.05947.x.

111. Pérez-Riverol Y, Bai J, Bandla C, García-Seisdedos D, Hewapathirana S, Kamatchinathan S, Kundu DJ, Prakash A, Frericks-Zipper A, Eisenacher M, Walzer M, Wang S, Brazma A, Vizcaíno JA. 2022. The PRIDE database resources in 2022: a hub for mass spectrometry-based proteomics evidences. Nucleic Acids Res 50:D543–D552. 10.1093/nar/gkab1038.

