## Supplementary Tables and Figures for "Adaptive laboratory evolution rewires *Pseudomonas putida* for resource-efficient acetate assimilation"

by

Nicolás Gurdo[a], Aparajitha Srinivasan[b,c], Tommaso Tagliani[a], Melanie Filbig[d], Nicolas T. Wirth[a], Josefin Johnsen[a], Garret W. O'Connell[a], Stefano Donati[a], Enrico Orsi[a], María V. G. Alván-Vargas[a], Yan Chen[b,c], Christopher J. Petzold[b,c], Matthew Blow[e], Thomas Eng[b,c], Till Tiso[d,f], Lars M. Blank[d,g], Adam Feist[a,b,h], Aindrila Mukhopadhyay[b,c,i], and Pablo I. Nikel[a]

[a] BRiGHT, Technical University of Denmark, Kgs. Lyngby, Denmark
[b] Joint BioEnergy Institute, Emeryville, CA, USA
[c] Biological Systems and Engineering Division, Lawrence Berkeley National Laboratory, Berkeley, CA, USA
[d] Institute of Applied Microbiology, RWTH Aachen University, Aachen, Germany
[e] U.S. Department of Energy Joint Genome Institute, Berkeley, CA, USA
[f] Systems Biotechnology, Technical Faculty, Bielefeld University, Bielefeld, Germany
[g] WSS Research Centre "CatalAix", Aachen, Germany
[h] Department of Bioengineering, University of California San Diego, La Jolla, CA, USA
[i] Environmental Genomics and Systems Biology Division, Lawrence Berkeley National Laboratory, Berkeley, CA, USA

### Table of contents

**Figure S1.** Transcript interaction network analysis of significantly **(A)** upregulated and **(B)** downregulated genes in the evolved strain compared with the parental strain.
**Figure S2.** Transcript interaction network analysis of significantly **(A)** upregulated and **(B)** downregulated genes in GacA$^{E197K}$ FabB$^{L77P}$ compared with the parental strain.
**Figure S3.** Protein interaction network analysis of significantly **(A)** upregulated and **(B)** downregulated proteins in the evolved strain compared with the parental strain.
**Figure S4.** Protein interaction network analysis of significantly **(A)** upregulated and **(B)** downregulated proteins in GacA$^{E197K}$ FabB$^{L77P}$ compared with the parental strain.
**Table S1.** Bacterial strains used in this study.
**Table S2.** Plasmids used in this study.
**Table S3.** Oligonucleotides used in this study.
**Table S4.** Carbon atom transitions used for $^{13}$C-metabolic flux analysis.

**Figure S1**. Transcript interaction network analysis of significantly **(A)** upregulated and **(B)** downregulated genes in the evolved strain compared with the parental strain.

## (A)

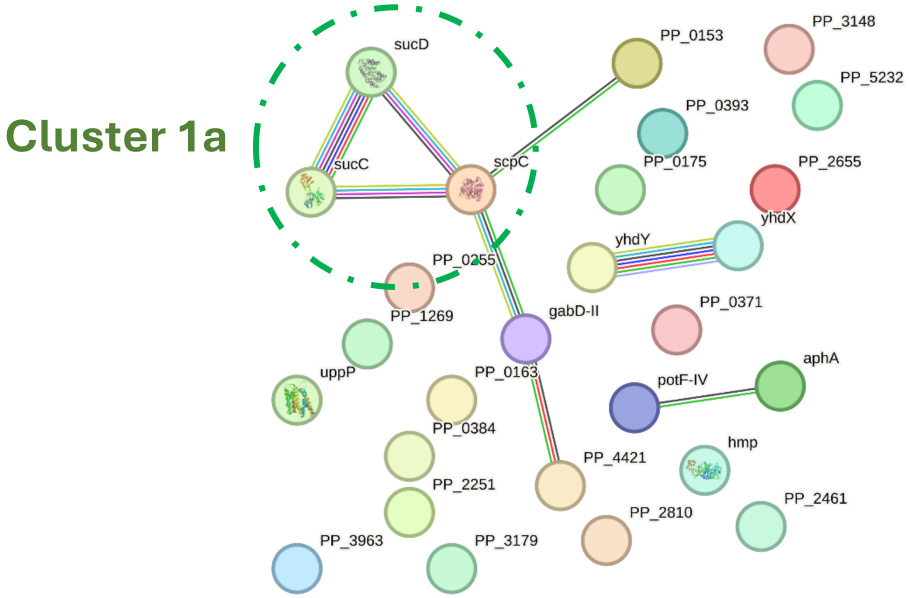

**Figure S1 (*cont.*).** Transcript interaction network analysis of significantly **(A)** upregulated and **(B)** downregulated genes in the evolved strain compared with the parental strain.

**(B)**

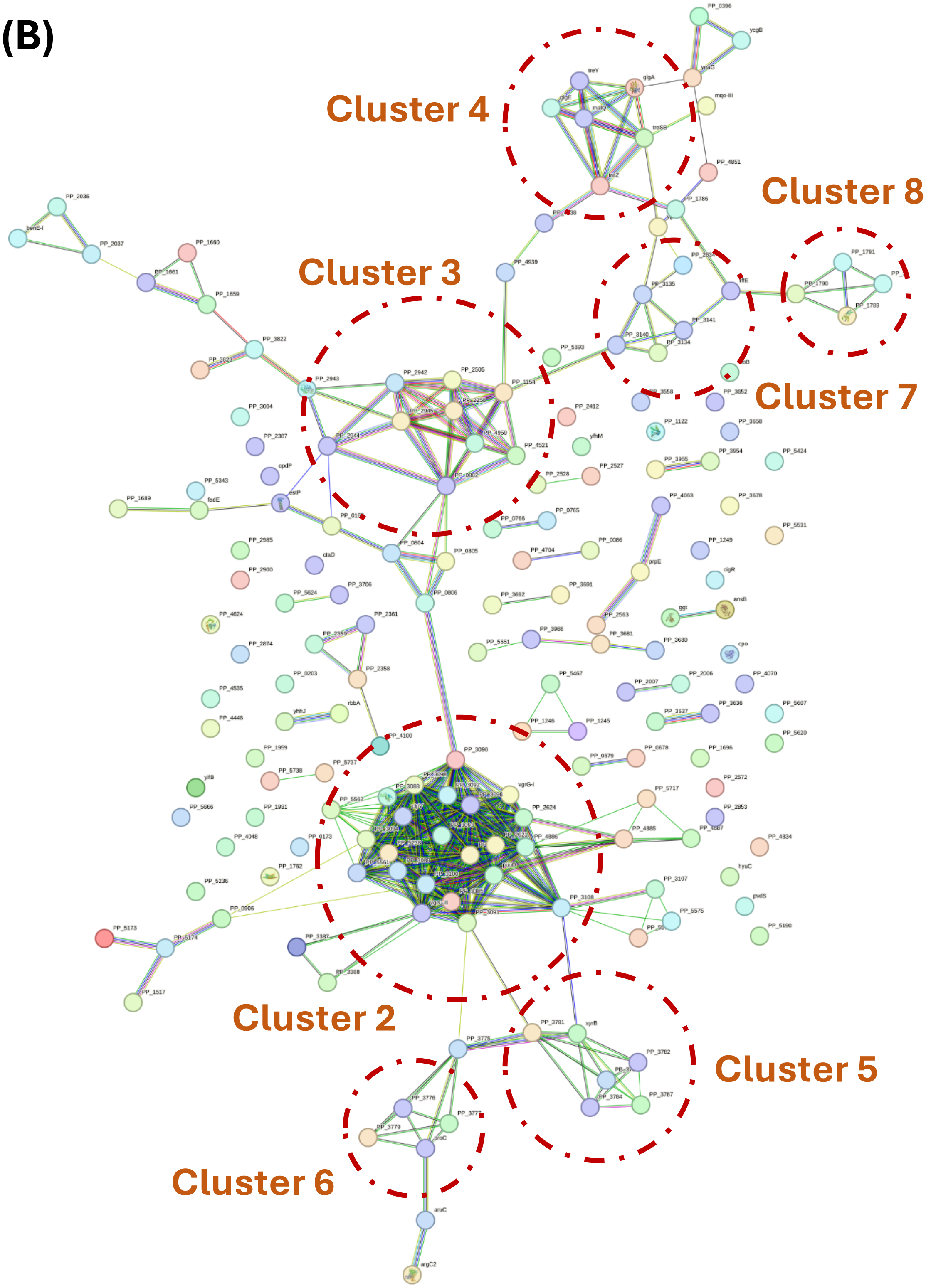

**Figure S2.** Transcript interaction network analysis of significantly **(A)** upregulated and **(B)** downregulated genes in GacA$^{E197K}$ FabB$^{L77P}$ compared with the parental strain.

# (A)

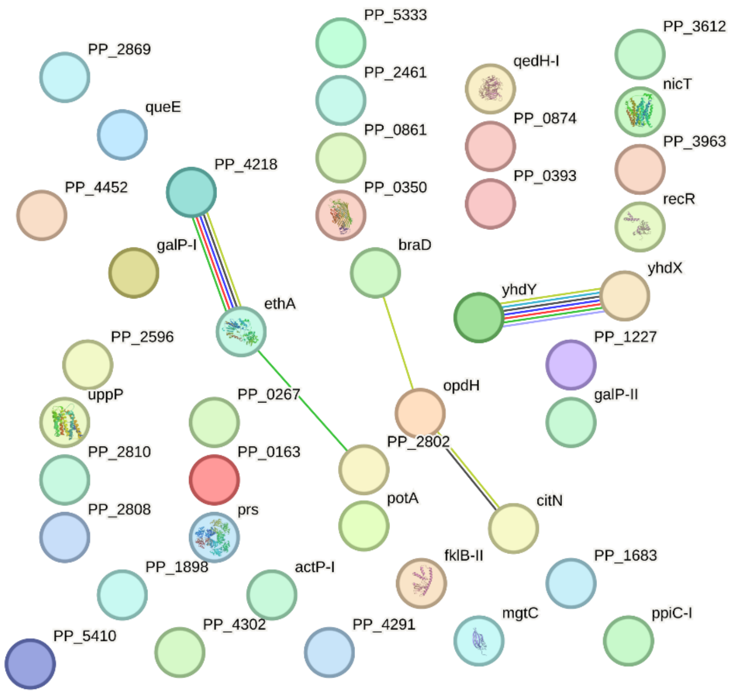

**Figure S2 (cont.).** Transcript interaction network analysis of significantly **(A)** upregulated and **(B)** downregulated genes in GacA$^{E197K}$ FabB$^{L77P}$ compared with the parental strain.

# (B)

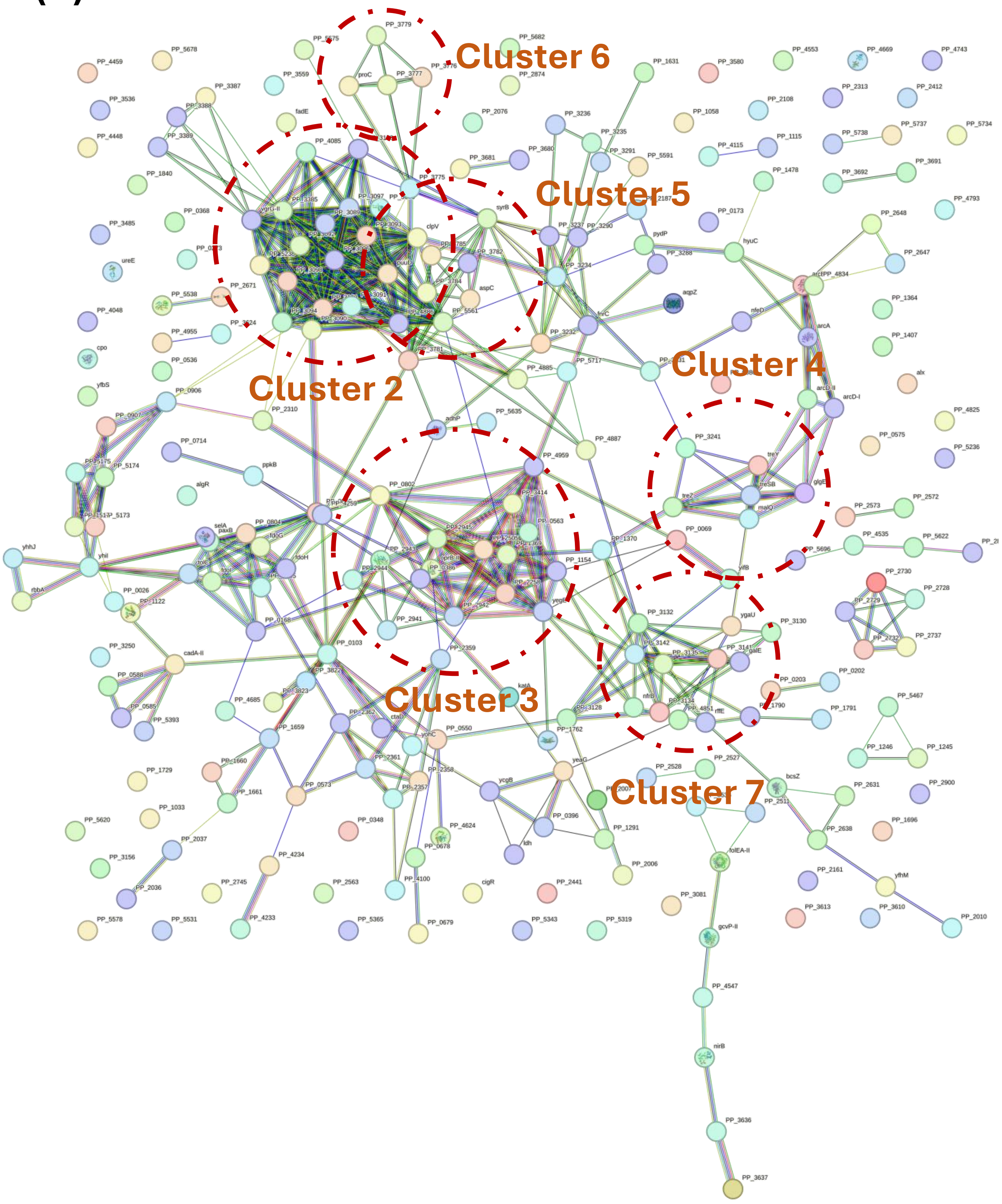

**Figure S3.** Protein interaction network analysis of significantly **(A)** upregulated and **(B)** downregulated proteins in the evolved strain compared with the parental strain.

## (A)

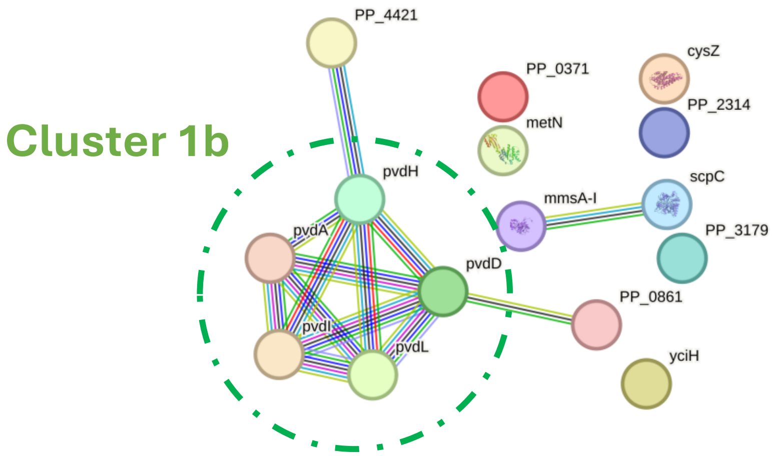

**Figure S3 (cont.).** Protein interaction network analysis of significantly **(A)** upregulated and **(B)** downregulated proteins in the evolved strain compared with the parental strain.

**(B)**

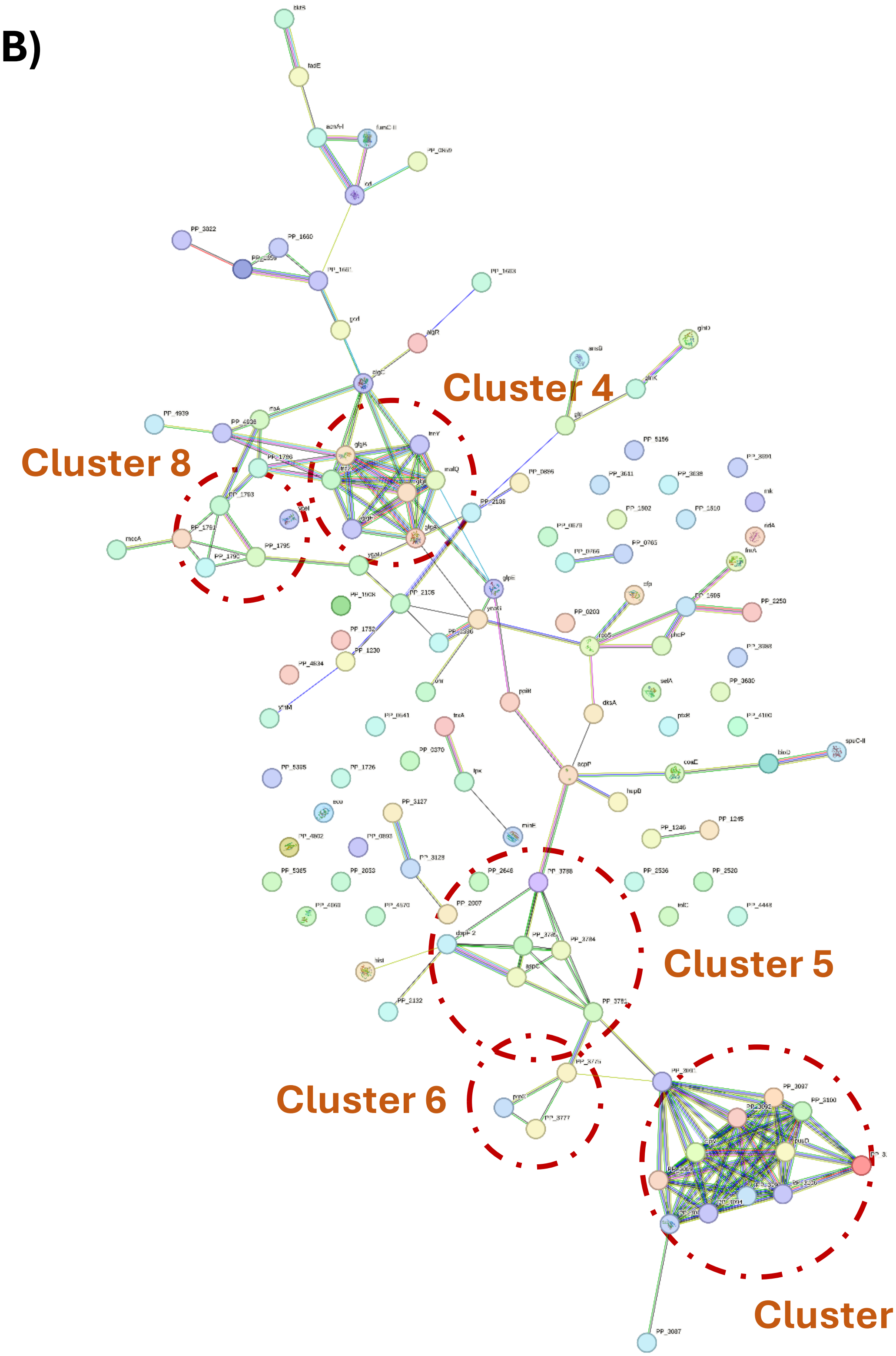

**Figure S4.** Protein interaction network analysis of significantly **(A)** upregulated and **(B)** downregulated proteins in GacA$^{E197K}$ FabB$^{L77P}$ compared with the parental strain.

## (A)

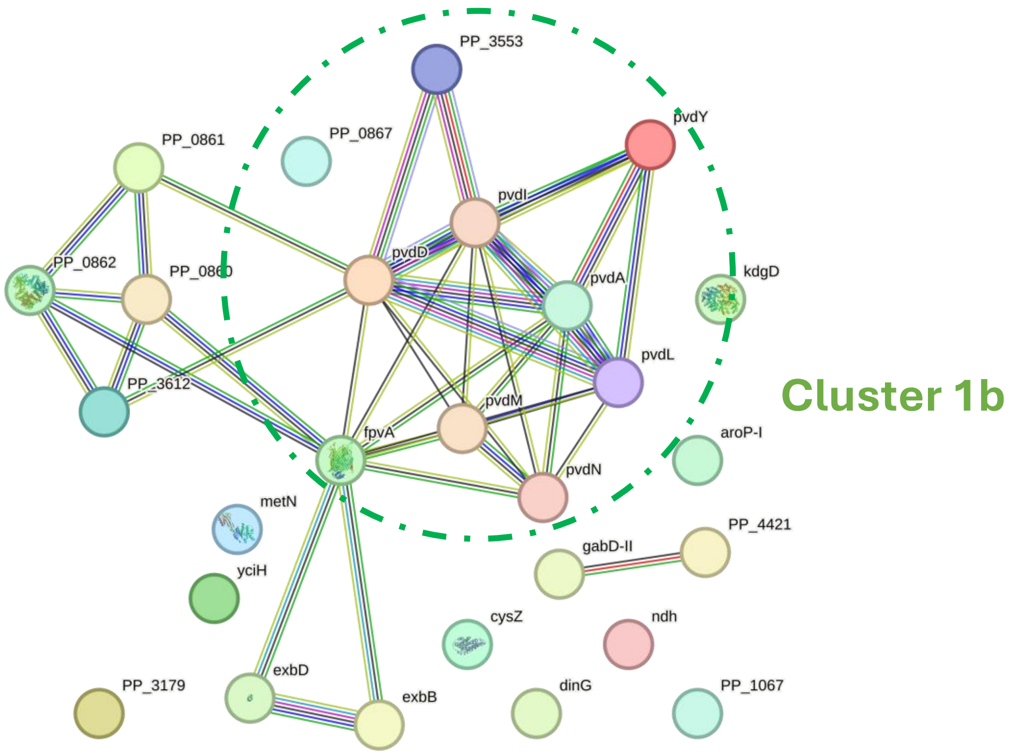

**Figure S4 (cont.).** Protein interaction network analysis of significantly **(A)** upregulated and **(B)** downregulated proteins in GacA$^{E197K}$ FabB$^{L77P}$ compared with the parental strain.

# (B)

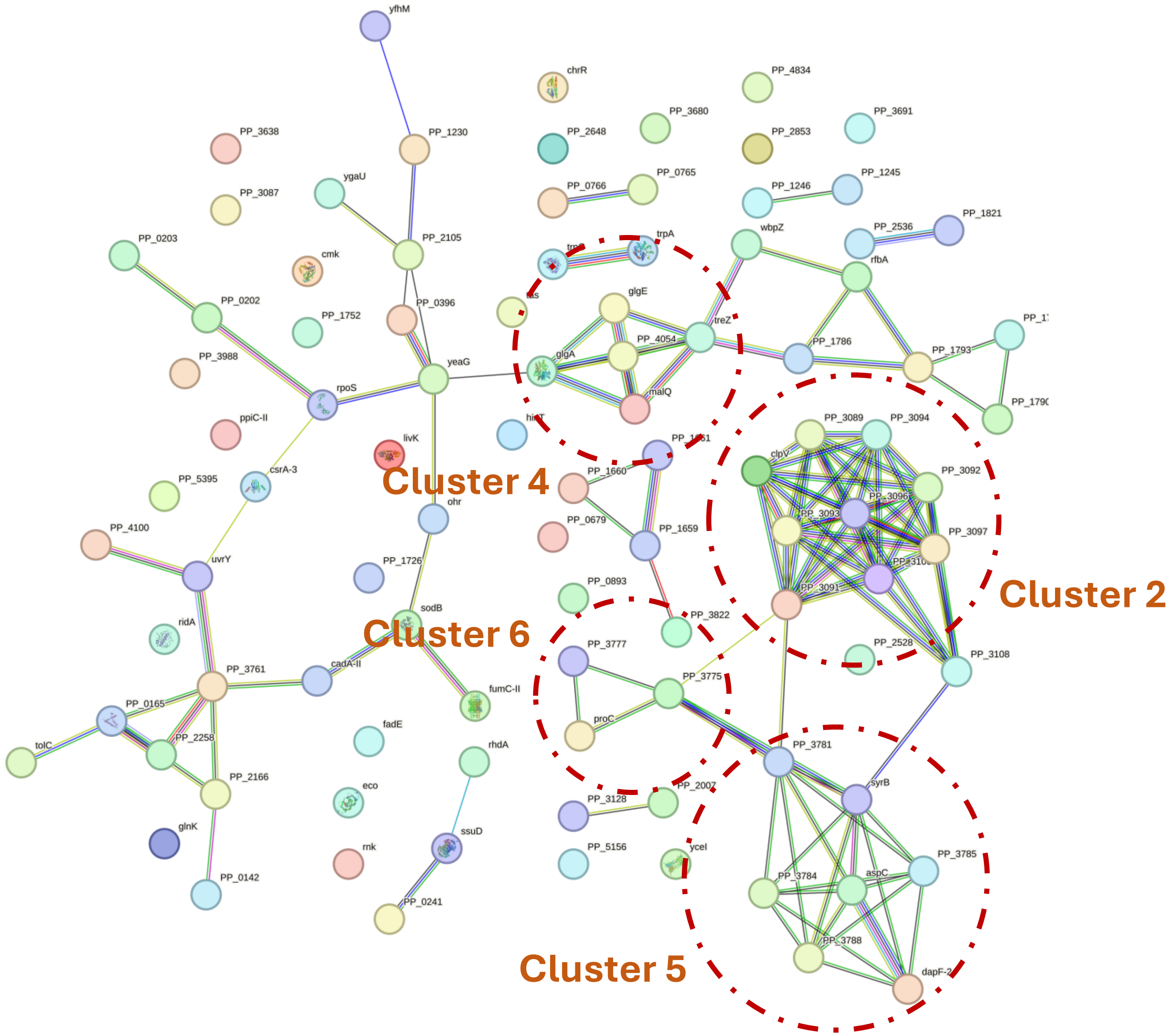

**Table S1. Bacterial strains used in this study.**

| Strain | Description[a] | Reference or source |
|---|---|---|
| ***Escherichia coli*** | | |
| DH5α λ*pir* | Cloning host; $F^-$ $\lambda^-$ *endA1 glnX44*(AS) *thiE1 recA1 relA1 spoT1 gyrA96*($Nal^R$) *rfbC1 deoR nupG* Φ80(*lacZ*ΔM15) Δ(*argF-lac*)*U169 hsdR17*($r_K^-$ $m_K^+$), λ*pir* lysogen | Platt et al. (1) |
| ***Pseudomonas putida*** | | |
| KT2440 | Wild-type strain, derived from *P. putida* mt-2 (2) cured of the TOL plasmid pWW0 | Bagdasarian et al. (3) |
| EM42 | Reduced-genome derivative of *P. putida* KT2440; Δ*PP4329-PP4397* (flagellar operon) Δ*PP3849-PP3920* (prophage I) Δ*PP3026-PP3066* (prophage II) Δ*PP2266-PP2297* (prophage III) Δ*PP1532-PP1586* (prophage IV) ΔTn*7* Δ*endA-1* Δ*endA-2* Δ*hsdRMS* ΔTn*4652* | Martínez-García et al. (4) |
| SEM1.4 | Reduced genome derivative of EM42; Δ*PP_5003-5008* (Δ*phaC1ZC2DFI*), Δ*PP_3161-3164* (Δ*benABCD*), Δ*PP_1408* (Δ*phaG*) | Ackermann et al. (5),Kozaeva et al. (6) |
| SEM1.4Evo | Evolved derivative of *P. putida* SEM1.4 with increased acetate tolerance | This work |
| SEM1.4 $HupB^{T49I}$ | *P. putida* SEM1.4 $HupB^{T49I}$ | This work |
| SEM1.4 $HupB^{A35V}$ | *P. putida* SEM1.4 $HupB^{A35V}$ | This work |
| SEM1.4 $FabB^{L77P}$ | *P. putida* SEM1.4 $FabB^{L77P}$ | This work |
| SEM1.4 $PP\_0371^{R277H}$ | *P. putida* SEM1.4 $PP\_0371^{R277H}$ | This work |
| SEM1.4 $GacA^{E197K}$ | *P. putida* SEM1.4 $GacA^{E197K}$ | This work |
| SEM1.4 $GacA^{E197K}$ $FabB^{L77P}$ | *P. putida* SEM1.4 $GacA^{E197K}$ $FabB^{L77P}$ | This work |

[a] Nal, nalidixic acid.

**Table S2. Plasmids used in this study.**

| Plasmid | Description[a] | Source |
|---|---|---|
| pSNW2 | Derivative of vector pGNW2 (7) with $P_{14g}$(BCD2)→*msfGFP*; $Km^R$ | Volke et al. (8) |
| pSNW::GacA$^{E197K}$ | Derivative of vector pSNW2 carrying homologous regions (HRs) to introduce the E197K substitution in *gacA*; $Km^R$ | This work |
| pSNW::HupB$^{T49I}$ | Derivative of vector pSNW2 carrying HRs to introduce the T49I substitution in *hupB*; $Km^R$ | This work |
| pSNW::HupB$^{A35V}$ | Derivative of vector pSNW2 carrying HRs to introduce the A35V substitution in *hupB*; $Km^R$ | This work |
| pSNW::PP_0371$^{R277H}$ | Derivative of vector pSNW2 carrying HRs to introduce the R277H substitution in *PP_0371*; $Km^R$ | This work |
| pSNW::FabB$^{L77P}$ | Derivative of vector pSNW2 carrying HRs to introduce the L77P substitution in *fabB*; $Km^R$ | This work |
| pQURE6·H | Conditionally-replicating vector; derivative of vector pJBSD1 carrying XylS/*Pm*→*I-SceI* and $P_{14g}$(BCD2)→ *mRFP*; $Gm^R$ | Volke et al. (8) |

[a] Gm, gentamicin; Km, kanamycin.

**Table S3. Oligonucleotides used in this study.**

| Primer | Nucleotide sequence (5'→3') | Use |
|---|---|---|
| **491_GacA$^{E197K}$_A_U_F** | agatcctGGCGGGCAGCCCGTAC | USER adaptors for mutagenesis of *gacA*$^{E197K}$ |
| **492_GacA$^{E197K}$_A_U_R** | AGTTTGACGTCGCTGGTGACC | |
| **493_GacA$^{E197K}$_B_U_F** | ACGTCAAACTGACCTTGCTGGC | |
| **494_ GacA$^{E197K}$_B_U_R** | aggtcgactGGCCGGGTGCGGTTGG | |
| **495_GacA$^{E197K}$_chk_F** | CTGACCAAGGGTGCAGGCCTTG | Sequencing *gacA* |
| **496_GacA$^{E197K}$_chk_R** | TGGCCTTGCCCACGTAAAGCAG | |
| **497_HupB$^{T49I}$_A_U_F** | agatcctGTGTCGGCGCTGACGCA | USER adaptors for mutagenesis of *hupB*$^{T49A}$ |
| **498_HupB$^{T49I}$_A_U_R** | AGAAGATACCAAAGCCAACCAGTACCAC | |
| **499_HupB$^{T49I}$_B_U_F** | ATCTTCTCGGTCAAGGAGCGC | |
| **500_HupB$^{T49I}$_B_U_R** | aggtcgactTGGCATCCTCAGCACCTTGCA | |
| **501_HupB$^{T49I}$_chk_F** | CCATGCCTCCCAGCACACGTTT | Sequencing *hupB* |
| **502_HupB$^{T49I}$_chk_R** | AAACGGCTGTACCACTGCGTCG | |
| **503_HupB$^{A35V}$_A_U_F** | agatcctGTGTCGGCGCTGACGCA | USER adaptors for mutagenesis of *hupB*$^{A35V}$ |
| **504_HupB$^{A35V}$_A_U_R** | ACGCCGGTGACGGATTCG | |
| **505_HupB$^{A35V}$_B_U_F** | ACCGGCGTCCTGAAGCAAG | |
| **506_HupB$^{A35V}$_B_U_R** | aggtcgactTGGCATCCTCAGCACCTTGCA | |
| **507_PP0371$^{R277H}$_A_U_F** | agatcctGAGCCCGGGTTGATCAACGC | USER adaptors for mutagenesis of *PP_0371*$^{R277H}$ |
| **508_PP0371$^{R277H}$_A_U_R** | AGGCCTGCTGGTGCTGGG | |
| **509_PP0371$^{R277H}$_B_U_F** | AGCAGGCCTCTGTGGAGCTG | |
| **510_PP0371$^{R277H}$_B_U_R** | aggtcgactGGCGGCGCTGGCG | |
| **511_PP0371$^{R277H}$_chk_F** | TGATGCTGGGCGAGGAGTTCCA | Sequencing *PP_0371* |
| **512_PP0371$^{R277H}$_chk_R** | CGGGGTTTTCTTTGCGTGCGTG | |
| **513_FabB$^{L77P}$_A_U_F** | agatcctGAAGACCTGCTGCGCTGCA | USER adaptors for mutagenesis of *fabB*$^{L77P}$ |
| **514_FabB$^{L77P}$_A_U_R** | ATGGCCGGGTAGGCGTAGG | |
| **515_FabB$^{L77P}$_B_U_F** | ACCCGGCCATGCAGGAC | |
| **516_FabB$^{L77P}$_B_U_R** | aggtcgactTACGCGATTCATCTGGGCGC | |
| **517_FabB$^{L77P}$_chk_F** | GTCGGCGTGTTCACCTCCACTG | Sequencing *fabB* |
| **518_FabB$^{L77P}$_chk_R** | AGGGTGTCCAGCGCTTCCATCT | |

**Table S4. Carbon atom transitions used for $^{13}$C-metabolic flux analysis.**

| Reaction # | Description and atom transitions |
|---|---|
| 1 | Ac.ext (ef) -> Ac (ef) |
| 2 | Ac (ef) + 2*ATP -> AcCoA (ef) |
| 3 | OAA (abcd) + AcCoA (ef) -> Cit (dcbfea) |
| 4 | Cit (abcdef) <-> ICit (abcdef) |
| 5 | ICit (abcdef) -> Suc (edcf) + Glyox (ab) |
| 6 | Glyox (ab) + AcCoA (cd) -> Mal (abdc) |
| 7 | ICit (abcdef) -> AKG (abcde) + CO2 (f) + NADPH |
| 8 | AKG (abcde) -> SucCoA (bcde) + CO2 (a) + NADH |
| 9 | SucCoA (abcd) <-> Suc (abcd) + ATP |
| 10 | Suc (abcd) -> Fum (abcd) + FADH2 |
| 11 | Fum (abcd) <-> Mal (abcd) |
| 12 | Mal (abcd) -> OAA (abcd) + FADH2 |
| 13 | Pyr (abc) + CO2 (d) + ATP -> OAA (abcd) |
| 14 | Pyr (abc) -> AcCoA (bc) + CO2 (a) + NADH |
| 15 | OAA (abcd) -> Pyr (abc) + CO2 (d) |
| 16 | PEP (abc) + CO2 (d) -> OAA (abcd) |
| 17 | OAA (abcd) + ATP -> PEP (abc) + CO2 (d) |
| 18 | Mal (abcd) -> Pyr (abc) + CO2 (d) + NADPH |
| 19 | G6P (abcdef) <-> F6P (abcdef) |
| 20 | FBP (abcdef) -> F6P (abcdef) |
| 21 | FBP (abcdef) <-> DHAP (cba) + GAP (def) |
| 22 | DHAP (abc) <-> GAP (abc) |
| 23 | GAP (abc) <-> 3PG (abc) + ATP + NADH |
| 24 | 3PG (abc) <-> PEP (abc) |
| 25 | PEP (abc) -> Pyr (abc) + ATP |
| 26 | G6P (abcdef) -> 6PG (abcdef) + NADPH |
| 27 | 6PG (abcdef) -> Ri5P (bcdef) + CO2 (a) + NADPH |
| 28 | Ri5P (abcde) <-> X5P (abcde) |
| 29 | Ri5P (abcde) <-> R5P (abcde) |
| 30 | X5P (abcde) <-> GAP (cde) + EC2 (ab) |

| | |
|---|---|
| 31 | F6P (abcdef) <-> E4P (cdef) + EC2 (ab) |
| 32 | S7P (abcdefg) <-> R5P (cdefg) + EC2 (ab) |
| 33 | F6P (abcdef) <-> GAP (def) + EC3 (abc) |
| 34 | S7P (abcdefg) <-> E4P (defg) + EC3 (abc) |
| 35 | 6PG (abcdef) -> Pyr (abc) + GAP (def) |
| 36 | AKG (abcde) + NADPH + NH3 -> Glu (abcde) |
| 37 | Glu (abcde) + ATP + NH3 -> Gln (abcde) |
| 38 | Glu (abcde) + ATP + 2*NADPH -> Pro (abcde) |
| 39 | Glu (abcde) + CO2 (f) + Gln (ghijk) + Asp (lmno) + AcCoA (pq) + 5*ATP + NADPH -> Arg (abcdef) + AKG (ghijk) + Fum (lmno) + Ac (pq) |
| 40 | OAA (abcd) + Glu (efghi) -> Asp (abcd) + AKG (efghi) |
| 41 | Asp (abcd) + 2*ATP + NH3 -> Asn (abcd) |
| 42 | Pyr (abc) + Glu (defgh) -> Ala (abc) + AKG (defgh) |
| 43 | 3PG (abc) + Glu (defgh) -> Ser (abc) + AKG (defgh) + NADH |
| 44 | Ser (abc) <-> Gly (ab) + MEETHF (c) |
| 45 | Gly (ab) <-> CO2 (a) + MEETHF (b) + NADH + NH3 |
| 46 | Thr (abcd) <-> Gly (ab) + AcCoA (cd) + NADH |
| 47 | Ser (abc) + AcCoA (de) + 3*ATP + 4*NADPH + SO4 -> Cys (abc) + Ac (de) |
| 48 | Asp (abcd) + Pyr (efg) + Glu (hijkl) + SucCoA (mnop) + ATP + 2*NADPH -> LL_DAP (abcdgfe) + AKG (hijkl) + Suc (mnop) |
| 49 | LL_DAP (abcdefg) -> Lys (abcdef) + CO2 (g) |
| 50 | Asp (abcd) + 2*ATP + 2*NADPH -> Thr (abcd) |
| 51 | Asp (abcd) + METHF (e) + Cys (fgh) + SucCoA (ijkl) + ATP + 2*NADPH -> Met (abcde) + Pyr (fgh) + Suc (ijkl) + NH3 |
| 52 | Pyr (abc) + Pyr (def) + Glu (ghijk) + NADPH -> Val (abcef) + CO2 (d) + AKG (ghijk) |
| 53 | AcCoA (ab) + Pyr (cde) + Pyr (fgh) + Glu (ijklm) + NADPH -> Leu (abdghe) + CO2 (c) + CO2 (f) + AKG (ijklm) + NADH |
| 54 | Thr (abcd) + Pyr (efg) + Glu (hijkl) + NADPH -> Ile (abfcdg) + CO2 (e) + AKG (hijkl) + NH3 |
| 55 | PEP (abc) + PEP (def) + E4P (ghij) + Glu (klmno) + ATP + NADPH -> Phe (abcefghij) + CO2 (d) + AKG (klmno) |
| 56 | PEP (abc) + PEP (def) + E4P (ghij) + Glu (klmno) + ATP + NADPH -> Tyr (abcefghij) + CO2 (d) + AKG (klmno) + NADH |

| | |
|---|---|
| 57 | Ser (abc) + R5P (defgh) + PEP (ijk) + E4P (lmno) + PEP (pqr) + Gln (stuvw) + 3*ATP + NADPH -> Trp (abcedklmnoj) + CO2 (i) + GAP (fgh) + Pyr (pqr) + Glu (stuvw) |
| 58 | R5P (abcde) + FTHF (f) + Gln (ghijk) + Asp (lmno) + 5*ATP -> His (edcbaf) + AKG (ghijk) + Fum (lmno) + 2*NADH |
| 59 | MEETHF (a) + NADH -> METHF (a) |
| 60 | MEETHF (a) -> FTHF (a) + NADPH |
| 61 | 0.174*G6P + 0.068*F6P + 0.107*GAP + 1.882*AcCoA + 0.431*Gly + 0.263*Pro + 0.598*Ala + 0.389*Val + 0.628*Leu + 0.244*Ile + 0.122*Met + 0.055*Cys + 0.191*Phe + 0.135*Tyr + 0.077*Trp + 0.126*His + 0.18*Lys + 0.354*Arg + 0.251*Gln + 0.158*Asn + 0.301*Glu + 0.284*Asp + 0.301*Ser + 0.256*Thr + 46.75*ATP -> Biomass |
| 62 | CO2_unlabeled (a) <-> CO2 (a) |
| 63 | NADH <-> NADPH |
| 64 | ATP -> ATP.maintenance |
| 65 | NADPH -> NADPH.maintenance |
| 66 | NADH + O2 -> 3*ATP |
| 67 | FADH2 + O2 -> 2*ATP |
| 68 | UQH2 + O2 -> 3*ATP |
| 69 | CO2 (a) -> CO2.ext (a) |
| 70 | NH3.ext -> NH3 |
| 71 | SO4.ext -> SO4 |
| 72 | O2.ext -> O2 |

### References


1. Platt R, Drescher C, Park SK, Phillips GJ. 2000. Genetic system for reversible integration of DNA constructs and *lacZ* gene fusions into the *Escherichia coli* chromosome. Plasmid 43:12–23. https://doi.org/10.1006/plas.1999.1433.
2. Worsey MJ, Williams PA. 1975. Metabolism of toluene and xylenes by *Pseudomonas putida* (*arvilla*) mt-2: evidence for a new function of the TOL plasmid. J Bacteriol 124:7–13. https://doi.org/10.1128/jb.124.1.7-13.1975.
3. Bagdasarian M, Lurz R, Rückert B, Franklin FCH, Bagdasarian MM, Frey J, Timmis KN. 1981. Specific purpose plasmid cloning vectors. II. Broad host range, high copy number, RSF1010-derived vectors, and a host-vector system for gene cloning in *Pseudomonas*. Gene 16:237–247. https://doi.org/10.1016/0378-1119(81)90080-9.
4. Martínez-García E, Nikel PI, Aparicio T, de Lorenzo V. 2014. *Pseudomonas* 2.0: genetic upgrading of *P. putida* KT2440 as an enhanced host for heterologous gene expression. Microb Cell Fact 13:159. https://doi.org/10.1186/s12934-014-0159-3.
5. Ackermann YS, de Witt J, Mezzina MP, Schroth C, Polen T, Nikel PI, Wynands B, Wierckx N. 2024. Bio-upcycling of even and uneven medium-chain-length diols and dicarboxylates to polyhydroxyalkanoates using engineered *Pseudomonas putida*. Microb Cell Fact 23:54. https://doi.org/10.1186/s12934-024-02310-7.
6. Kozaeva E, Volkova S, Matos MRA, Mezzina MP, Wulff T, Volke DC, Nielsen LK, Nikel PI. 2021. Model-guided dynamic control of essential metabolic nodes boosts acetyl-coenzyme A–dependent bioproduction in rewired *Pseudomonas putida*. Metab Eng 67:373–386. https://doi.org/10.1016/j.ymben.2021.07.014.
7. Wirth NT, Kozaeva E, Nikel PI. 2020. Accelerated genome engineering of *Pseudomonas putida* by I-*Sce*I—mediated recombination and CRISPR-Cas9 counterselection. Microb Biotechnol 13:233–249. https://doi.org/10.1111/1751-7915.13396.
8. Volke DC, Friis L, Wirth NT, Turlin J, Nikel PI. 2020. Synthetic control of plasmid replication enables target- and self-curing of vectors and expedites genome engineering of *Pseudomonas putida*. Metab Eng Commun 10:e00126. https://doi.org/10.1016/j.mec.2020.e00126.
